# Social-Cognitive Dysregulation Model of Misophonia: Perspective from a Behavioural Study

**DOI:** 10.64898/2026.09.03.749152

**Authors:** Mercede Erfanian, Sukhbinder Kumar

## Abstract

Misophonia is increasingly conceptualized as more than a disorder of sound tolerance, with trigger over-reactivity shaped by the social meaning of sounds, inferred intentions, and representations of others’ actions. We tested a social-cognitive dysregulation model of misophonia in (N = 341) adults using behavioural measures of Theory of Mind and emotion recognition, alongside measures of reflective functioning, empathy, alexithymia, and mimicry. Dimensional associations with misophonia severity and its five different dimensions were examined while accounting for age, sex, sound sensitivity, and anxiety/depressive symptoms. Increased misophonia severity was associated with less accurate and slower mental-state inference and emotion recognition. ToM accuracy effects were evident for more complex, cognitive, and affective mentalizing, but not for simpler mentalizing or physical control judgments, while emotion-recognition accuracy differences emerged for positive but not negative stimuli. Greater severity was also characterized by reduced certainty and greater uncertainty about mental states, greater difficulty identifying one’s own feelings, and elevated alexithymia, whereas global self-reported empathy was largely preserved. Misophonia severity further predicted a greater propensity to mimic trigger-producing actions or sounds; 41% of participants exceeding the S-Five clinical cutoff (≥87) endorsed mimicry, which was particularly associated with a subjective restoration of control. Findings remained robust following influential-case sensitivity analyses. These results reveal a selective disturbance in self–other representation spanning mentalizing, emotion decoding, emotional self-representation, and embodied regulatory processes. They position misophonia within a broader social-cognitive framework in which auditory-affective reactivity may intersect with altered inferential and sensorimotor processing, while stopping short of causal inference.

## 1. Introduction

Misophonia is a disorder phenotypically characterized by intense and disproportionate aversive responses to specific everyday sounds, most commonly human-generated orofacial sounds such as chewing, breathing, or swallowing, although its triggers extend well beyond this category of auditory triggers (Black et al., 2025; Brout et al., 2018; Swedo et al., 2022; Vitoratou et al., 2023; S. Vitoratou et al., 2021).

Since the term was introduced, research has documented a wide range of behavioural, emotional, physiological, and cognitive characteristics both as function of triggers exposure and in their absence, alongside frequent co-occurrence with psychiatric and audiological conditions that share some of its symptoms (Aazh et al., 2022a; Daniels et al., 2020; de Gee et al., 2026; Eijsker et al., 2019; Erfanian et al., 2018a; Erfanian et al., 2019; Erfanian et al., 2018b; Kumar et al., 2017; Oszczapinska et al., 2025; Rouw et al., 2018; Schroder et al., 2019; S. Vitoratou et al., 2021). Its symptoms vary considerably across individuals, its triggers are highly idiosyncratic, and its phenomenology overlaps with multiple clinical conditions without being fully explained by any of them. Early research, shaped largely by a bottom-up perspective, focused on the acoustic properties of trigger sounds and the intense emotional reactions they elicited (Clonan et al., 2025; Kirby et al., 2025; Schroder et al., 2014). More recent behavioural and neurobiological evidence, however, suggests that these responses cannot be understood merely in terms of abnormal sound processing. Instead, misophonia may arise from a dynamic interaction characterized by context based differential weighting of sensory input and higher-order processes that assign meaning, salience, intention, and social significance to that input (Banker et al., 2022; Berger et al., 2024; Black et al., 2025; Du et al., 2026; Eijsker et al., 2019; Hanna et al., 2026; Hansen et al., 2024; Humolli et al., 2025; Kumar et al., 2021; Kumar et al., 2017; Savard et al., 2025). This emerging perspective invites a far-reaching conceptualisation of misophonia, not only as a disorder of sound tolerance or sensory-emotional reactivity, but also as a disorder that may involve fundamental aspects of social cognition.

Misophonic over-reactivity cannot be reduced to the acoustic properties of the eliciting stimulus (i.e., loudness). Rather, response magnitude appears to be modulated by the identity of the sound producer, causal attributions regarding the sound source, and the interpersonal meaning assigned to the sound-producing act (Berger et al., 2024; Jastreboff et al., 2014; McGeoch et al., 2020; Ozuer et al., 2025; Rosenthal et al., 2026; Siepsiak et al., 2023). Acoustically similar sounds can elicit substantially different affective responses across social contexts, with greater aversion reported when sounds are produced by negatively evaluated individuals; mutually, the production of a trigger sound can itself alter social evaluations of the person responsible (Humolli et al., 2025; Siepsiak et al., 2023). Experimental findings suggest that otherwise innocuous orofacial sounds may acquire maladaptive meanings related to intrusion, offence, boundary violation, and diminished autonomy (Ozuer et al., 2025). Misophonic triggers may therefore operate not simply as aversive sensory inputs, but as socially encoded cues whose pathological salience emerges through higher-order appraisal of another person actions and intentions (Berger et al., 2024; Humolli et al., 2025; Ozuer et al., 2025).

Interpreting such signals requires social cognition, the set of processes through which individuals represent, infer, and respond to the internal states of others (Frith et al., 2007). These processes include Theory of Mind (ToM), or mentalizing, the capacity to infer beliefs, intentions, and perspectives, as well as cognitive and affective components of empathy (Preckel et al., 2018; Schurz et al., 2021). Although these capacities interact, they rely on partly distinguishable processes, ranging from abstract inference about another person mental state to affective and embodied representations of their experience (Schurz et al., 2021). If misophonic reactions are predominantly driven by the interpretation of another person behaviour and intention, then individual differences in mental-state inference and emotional understanding may facilitate our understanding on why socially generated sounds acquire such disproportionate salience.

Understanding others, however, is somewhat grounded in the representation of one own internal states (Luyten et al., 2015). Alexithymia, a dimensional difficulty in identifying and describing feelings (Luminet et al., 2021), may therefore be particularly relevant. Difficulties differentiating emotional states from bodily arousal could influence how intense sensory–affective reactions are interpreted and how the emotions of others are recognised (Demers et al., 2015; Moriguchi et al., 2006). In addition, alexithymia is consistently associated with a deficit of empathy (Banzhaf et al., 2018) and heterogeneous patterns involving impaired emotion processing, preserved mental-state recognition, and heightened self-oriented distress in response to others’ emotions (Luminet et al., 2021; Nam et al., 2020). This distinction may be of particular explanatory value for understanding misophonia, a) because misophonia is tied to altered processing in brain areas implicated in interoception (Kumar et al., 2017) and b) where intense internal arousal and higher-order evaluation may jointly shape responses to socially generated trigger events (Savard et al., 2022; Swedo et al., 2022).

Embodied self–other processes may provide a complementary pathway. Neuroimaging evidence indicates increased engagement and connectivity of orofacial motor cortex when individuals with misophonia perceive trigger sounds, supporting the notion that the actions of others may be excessively represented within the listener motor system (Kumar et al., 2021). Relatedly, many individuals with misophonia report mimicking trigger-producing actions or sounds, and the frequency of this behaviour is associated with symptom severity and trigger type (Ash et al., 2023; Rouw et al., 2018). Since mimicry can contribute to emotional understanding and affiliation, while remaining sensitive to social appraisal and context, altered mimicry may reflect both an attempt to regulate misophonic distress and a broader alteration in embodied self–other processing (Kastendieck et al., 2021; Rauchbauer et al., 2020).

All told, misophonia may reflect alterations in ‘self–other emotional and cognitive processing’, rather than an isolated abnormality of auditory perception. From this model, heightened misophonia severity should be associated with a) reduced efficiency in interpreting the mental and emotional states of others, b) greater difficulty identifying and representing one’s own emotional states, and c) altered mimicry. This model does not predict a generalized social-cognitive impairment. Instead, misophonia may be characterised by a selective profile across cognitive and affective mentalizing, emotion recognition, and empathic responsiveness.

So, the present study tested this integrative model by I) examining associations between misophonia severity and ToM, II) cognitive and affective empathy, III) alexithymia, and IV) mimicry. We hypothesised that greater misophonia severity would be associated with poorer and/or slower performance on ToM, affective and cognitive empathy tasks, higher alexithymia and altered mimicry. We further examined whether these associations were specific to particular components of social cognition and whether they remained after accounting for age, sex, anxiety, depression, and sound sensitivity. Given the cross-sectional design, these variables were treated as interrelated components of a conceptual model rather than as established causal mechanisms.

## 2. Method

The study was approved by ESSCA School of Management ethics committee (Avis. 21.2026). The study was preregistered on Open Science Framework (OSF) [https://osf.io/vjkfx] before the data collection. All raw, processed data and MATLAB code written for this study are openly available at [https://osf.io/mdnrj].

### 2.1. Participants

The sample comprised 341 participants (Mean_age_ = 34.68 years, SD = 10.02). Age and sex data were available for 329 participants, of whom 167 identified as male (50.76%) and 162 as female (49.24%). Participants were recruited through Prolific (www.prolific.com), an online research-participant platform (Figure 1).

**Figure 1.**
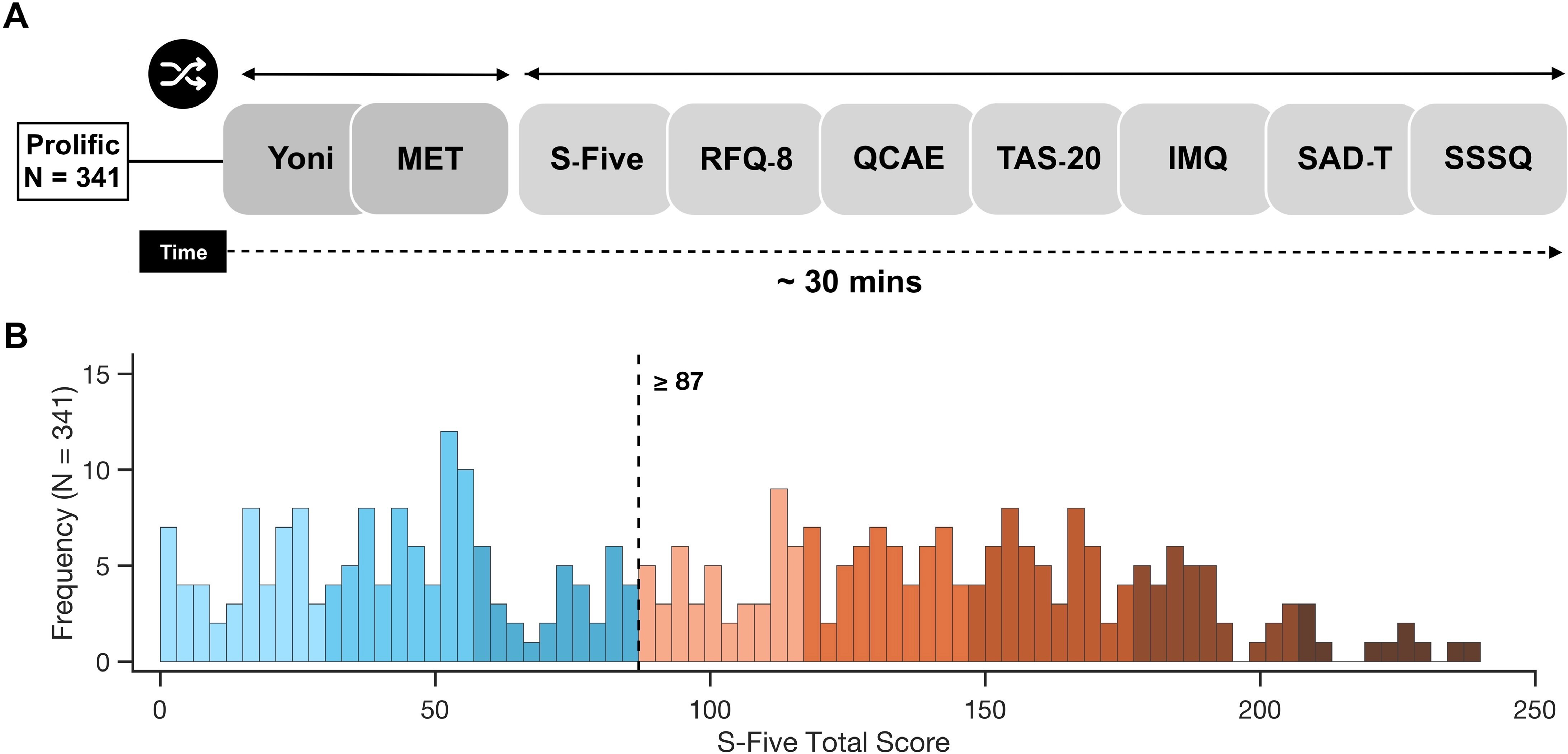
Study procedure and distribution of misophonia severity (N = 341) Overview of the online study. Participants completed the Yoni task and Multifaceted Empathy Task (MET), and in the task block, and the Selective Sound Sensitivity Syndrome Scale (S-Five), Reflective Functioning Questionnaire (RFQ-8), Questionnaire Of Cognitive And Affective Empathy (QCAE), Toronto Alexithymia Scale (TAS-20), Iowa Mimicry Questionnaire (IMQ), Sound Sensitivity Symptoms Questionnaire (SSSQ), and Screening for Anxiety and Depression in Tinnitus–Hyperacusis–Misophonia (SAD-T) in the questionnaire block; order was randomized within blocks. The battery took approximately 30 min. (B) Distribution of S-Five Total scores in the full sample (N = 341). The dashed line marks the prespecified clinical cutoff of ≥87(Vitoratou et al., 2023).

Following recommendations for transparent sample-size justification (Lakens, 2022), we aimed to recruit the largest practically feasible sample given the specificity of the target population. An a priori power analysis was conducted in R using the ‘pwrss’ package for the primary hierarchical multiple-regression model. Assuming a small-to-medium incremental effect of f^2^ = .063, a = .05, and 95% power, the minimum required sample was 210 participants. This effect was selected as the smallest effect of theoretical relevance, prioritizing substantive justification and empirical precedent over conventional benchmarks (Correll et al., 2020), while accounting for potential inflation in published effect estimates (Anderson et al., 2017). Black et al. (2025) reported robust effects using similar behavioural tasks in a sample of 140 participants. The sample is therefore considered sufficient to detect theoretically meaningful effects.

### 2.2. Materials

#### 2.2.1. Behavioural Tasks

##### 2.2.1.1. Yoni Task

ToM was assessed using the 48-item short form of the Yoni task (Isernia et al., 2023; Shamay-Tsoory et al., 2007), a computerised measure designed to capture the multidimensional and hierarchical architecture of mentalizing. The task differentiates cognitive ToM, involving inferences about beliefs, intentions, and thoughts, from affective ToM, involving inferences about emotional states as well as physical control. Physical control assesses basic visuospatial processing and physical judgments (e.g., identifying the object closest to Yoni) without requiring mental-state attribution. It therefore functions primarily as a non-mentalizing control condition, helping to determine whether observed effects are specific to mental-state processing rather than reflecting more general visuospatial or task-related difficulties. It also further distinguishes first-order mentalizing, which requires representing Yoni immediate mental state, from second-order mentalizing, which requires inferring Yoni representation of another person’s mental state. These two components may include different trials of cognitive, affective ToM and physical control. On each trial, Yoni appeared at the centre of the screen, surrounded by four candidate objects or characters. Participants were presented with a brief prompt concerning what Yoni was thinking, intending, liking, or feeling and selected the response option that best matched his mental state for an instance “Yoni is thinking of …”. Correct responding required the integration of verbal information with gaze direction, facial expression, and contextual cues (See procedure).

The Yoni-48 total-score reliability was strong across multiple indices (ω = .86, α = .90, Guttman’s λ₂ = .91, λ₆ = .95), with high split-half reliability (Spearman–Brown ρ = .90). Item-level analyses using classical test theory and Rasch modelling further indicated particularly strong discrimination for second-order cognitive and affective mentalizing items (Isernia et al., 2023).

##### 2.2.1.2. Multifaceted Empathy Test (MET)

Task-based empathy was assessed using the English-language MET (Dziobek et al., 2008; Foell et al., 2018), a computerized paradigm distinguishing cognitive empathy, the ability to identify another person’s emotional state, from affective empathy, the emotional response elicited by that state. The MET comprises 40 naturalistic photographs depicting individuals in emotionally salient situations of either positive or negative valence. Essentially, it evaluates empathy by selecting the most appropriate mental-state descriptor from four alternatives. In the present study, accuracy and response time (RT) were examined overall and separately for positive- and negative-valence stimuli adapted from valence-specific MET analyses (Hurlemann et al., 2010; Wingenfeld et al., 2014) (See procedure).

Foell et al. (2018) reported high internal consistency for affective empathy (α = .94), whereas reliability was lower for cognitive-empathy accuracy (KR-20 = .49), increasing only marginally after item refinement (KR-20 = .51).

#### 2.2.2. Questionnaires

##### 2.2.2.1. Selective Sound Sensitivity Syndrome Scale (S-Five)

We used 25-item S-Five (Vitoratou et al., 2021) to quantify five symptom dimensions including externalising, internalising, impact, outbursts, and threat, alongside a total severity score. Items are rated on an 11-point scale from 0 (not at all true) to 10 (completely true), yielding total scores from 0 to 250, with higher scores indicating greater symptom severity. A total score of 87 or above has been proposed as indicative of clinically significant misophonia on the basis of receiver operating characteristic (ROC) analysis. The S-Five has demonstrated good internal consistency across its subscales (α ≥ .83), strong test–retest reliability (ICC ≥ .86), and satisfactory convergent and discriminant validity (Vitoratou et al., 2023). In the present sample, the total scale showed excellent internal consistency (α = .972, ω = .972; N = 338).

##### 2.2.2.2. Reflective Functioning Questionnaire (RFQ-8)

Alongside Yoni task we also measured mentalizing capacity using the eight-item RFQ-8 (Fonagy et al., 2016), a brief self-report measure of the ability to understand oneself and others in terms of intentional mental states, including emotions, beliefs, desires, and goals. Items are rated on a 7-point scale from 1 (strongly disagree) to 7 (strongly agree) and recoded to derive two distinct six-item dimensions: certainty about mental states (RFQ-C, hypermentalizing) and uncertainty about mental states (RFQ-U, hypomentalizing). Higher RFQ-C scores indicate greater confidence in interpreting mental states, whereas higher RFQ-U scores reflect greater ambiguity and difficulty in understanding one’s own and others’ internal experiences. Given the RFQ-8 does not yield a psychometrically valid global score, the two dimensions were analysed separately. Internal consistency in this study was good for RFQ-C (α = .843, ω = .847) and acceptable for RFQ-U (α = .764, ω = .768; N = 341).

##### 2.2.2.3. Questionnaire of Cognitive and Affective Empathy (QCAE)

Self-reported empathy was evaluated by using the 31-item QCAE (Reniers et al., 2011), which distinguishes the capacity to represent another person’s emotional state from the tendency to respond affectively to that state. The measure comprises two higher-order domains. Cognitive empathy which includes perspective taking (10 items) and online simulation (9 items), whereas affective empathy comprises emotion contagion (4 items), peripheral responsivity (4 items), and proximal responsivity (4 items). Participants rated each statement on a 4-point scale from 1 (strongly disagree) to 4 (strongly agree). Four negatively keyed items were reverse-scored before subscale scores were summed; higher scores indicate greater self-reported empathy.

Its internal consistency ranged from acceptable to good across subscales (α = .65 - .85), and the second-order model showed satisfactory fit (RMSEA = .077, CFI = .925, TLI = .908, SRMR = .042). Convergent validity was underpinned by strong associations with corresponding dimensions of the Basic Empathy Scale (cognitive empathy: r = .62; affective empathy: r = .76) (Reniers et al., 2011). The full QCAE demonstrated good internal consistency (α = .882, ω = .883; N = 341) in this study.

##### 2.2.2.4. Toronto Alexithymia Scale (TAS-20)

The TAS-20 was applied to measure Alexithymia (Bagby et al., 1994). TAS is a self-report measure of difficulties in identifying and articulating emotional states and a tendency toward externally oriented cognition. The scale comprises three dimensions, difficulty identifying feelings (DIF) (7 items), difficulty describing feelings (DDF) (5 items), and externally oriented thinking (EOT) (8 items). Items are rated on a 5-point scale from 1 (strongly disagree) to 5 (strongly agree); items 4, 5, 10, 18, and 19 are reverse-scored. Total scores range from 20 to 100, with higher scores indicating greater alexithymic traits.

The original validation supported a stable three-factor structure across nonclinical and psychiatric samples, with acceptable internal consistency for the total scale (α = .81) and subscales (α = .66 - .78), alongside good 3-week test–retest reliability (r = .77) (Bagby et al., 1994). In the present sample, the total scale showed modest to acceptable internal consistency (α = .657, ω = .711; N = 341).

##### 2.2.2.5. Iowa Mimicry Questionnaire (IMQ)

IMQ including a set of five questions, was used. It is designed to assess prevalence of mimicry and its effect on perceived distress in misophonia (Ash et al., 2023). The initial screening item asked whether participants ever engaged in such mimicry, with responses coded as 1 (yes) and 0 (no). Participants who endorsed mimicry completed four conditional follow-up items assessing whether the behaviour was spontaneous and uncontrollable, deliberate and voluntarily initiated, associated with relief from distress, and accompanied by a restored sense of control. Responses were recorded on a 5-point frequency scale from 0 (never) to 4 (always); participants who did not endorse mimicry were coded as ‘no mimicry’.

Mimicry endorsement was analysed as a binary outcome, whereas the four follow-up items were examined separately as distinct phenomenological characteristics of mimicry and are not designed to be summed into one total score. No composite score or internal consistency estimate was calculated.

##### 2.2.2.6. Sound Sensitivity Symptoms Questionnaire (SSSQ)

We measured sound-intolerance symptoms (i.e., hyperacusis) the six-item SSSQ (Aazh et al., 2024b), which captures features of tinnitus, hyperacusis and misophonia. Participants indicated the frequency of each symptom over the preceding 14 days using a 4-point scale from 0 to 3 (e.g. 0–1, 2–6, 7–10 and 11–14llldays), yielding total scores from 0 to 18; scores of 4 or above indicate clinically relevant sound sensitivity. The SSSQ was included to account for sound intolerance, particularly hyperacusis, which frequently co-occurs with misophonia (Aazh et al., 2022a; Andermane et al., 2023). The original five-item version demonstrated good internal consistency (α = .87) and satisfactory convergent and construct validity. Preliminary psychometric evaluation of the six-item version indicated good internal consistency (α = .80, ω = .80) and test–retest reliability (r = .81). In the present sample, internal consistency was excellent for the full scale (α = .909, ω = .910; N = 341).

##### 2.2.2.7. Screening for Anxiety and Depression in Tinnitus–Hyperacusis–Misophonia (SAD-T)

A four-item SAD-T (Aazh et al., 2022b), a brief measure validated in populations with auditory conditions. Participants rated how frequently they were troubled by symptoms such as feeling nervous or on edge and feeling down or hopeless presented with similar response options as the SSSQ (e.g. 0–1 days). Scores of 4 or above indicate clinically relevant anxiety or depressive symptomatology. The SAD-T has demonstrated strong internal consistency (α = .91) and corrected item–total correlations ranging from .76 to .84 (Aazh et al., 2024a). Internal consistency was good in the present sample (α = .882, ω = .883; N = 341).

### 2.3. Procedure

Participants were recruited through Prolific. Prespecified eligibility criteria were age 18–60 years, no self-reported hearing loss or hearing difficulties, no cognitive impairment or dementia, normal or corrected-to-normal vision, and being native or bilingual English speaker. Eligible participants viewed a study advertisement outlining the procedures and compensation of approximately £4 (∼€4.7 or $5.33), corresponding to an estimated hourly rate of £9.68 (∼€10.92 or $13.02).

All participants accessed the study through Gorilla Experiment Builder (www.gorilla.sc). At entry, they indicated whether they identified as having misophonia, had received a diagnosis, were unfamiliar with the condition or no misophonia. Prolific participants then entered their Prolific ID. Participants reviewed the information sheet and provided informed consent before proceeding. The experimental battery began with the Yoni task and the MET, which were always administered before the questionnaire battery but were randomised in order (task block). This sequencing was intended to minimise fatigue-related effects on behavioural accuracy and RT. Participants subsequently completed the S-Five, RFQ-8, QCAE, TAS-20, IMQ, SSSQ, and SAD-T, with questionnaire order randomised across participants (questionnaire block). All trials per task and questions per questionnaire were also fully randomized (Figure 1A). Standardised instructions preceded each measure. Five attention checks were embedded throughout the battery, one in each behavioural task and three across the questionnaire measures.

The study required approximately 30 minutes, with a median completion time of approximately 24 minutes. Prolific participants who completed the battery adequately received a completion code and were redirected to Prolific for payment. Participants failing two or more attention checks would have received a separate code and been asked to return their submission without compensation; no participant met this exclusion criterion. Demographic information provided through Prolific including age and gender, was linked to participants Gorilla data using their Prolific ID. These variables were used to characterise the sample and were included in the primary analyses as covariates.

A total of 717 participants were recruited. Participants were excluded for incomplete behavioural-task data, multiple participation attempts, or incomplete questionnaire data (N = 376), yielding a final analytic sample of N = 341, including 146 below and 195 above cutoff threshold (≥87) (Vitoratou et al., 2023) (Figure 1B). No participant failed two or more of the five embedded attention checks.

### 2.4. Study Variables

#### 2.4.1. Predictor variables

The primary predictor was dimensional misophonia severity, indexed by the S-Five total score. Secondary analyses examined the five S-Five dimensions, externalising, internalising, impact, outburst, and threat as separate predictors.

#### 2.4.2. Outcome variables

Outcomes included Yoni task accuracy (%) and RT, examined overall and across cognitive, affective, first-order, and second-order conditions; MET accuracy (%) and RT for overall and positive- and negative-valence indices; RFQ-8 certainty and uncertainty; QCAE cognitive and affective empathy, together with perspective taking, online simulation, emotion contagion, peripheral responsivity, and proximal responsivity as individual constructs; TAS-20 total alexithymia, DIF, DDF, and EOT; and IMQ mimicry endorsement, spontaneous mimicry, deliberate mimicry, distress relief, and perceived control.

#### 2.4.3. Covariates (control)

Age, sex, sound sensitivity, anxiety and depressive symptoms were included as covariates to evaluate the specificity of associations between misophonia severity and social-cognitive functioning.

Hyperacusis-related symptoms were controlled since participants with clinically significant misophonia showed elevated sound-sensitivity scores even after excluding the misophonia-specific item, consistent with the frequent co-occurrence of misophonia with hyperacusis and tinnitus (Aazh et al., 2022a). Nevertheless, misophonia can occur in the absence of identifiable audiological abnormalities (Swedo et al., 2022). Anxiety and depressive symptoms were controlled because both are prevalent among individuals with misophonia and could contribute independently to variation in emotional and social-cognitive functioning (Aazh et al., 2022a; Brout et al., 2018; Erfanian et al., 2018a; Erfanian et al., 2019; Erfanian et al., 2018b; Rouw et al., 2018).

At the same time, there has been ongoing debate regarding the extent to which misophonia may be accounted for by anxiety-related symptomatology (Rosenthal et al., 2026), although existing evidence suggests that anxiety explains only a relatively small proportion of variance in misophonia severity (Norris et al., 2022). Nevertheless, to examine this issue directly, we additionally repeated the analyses using S-Five severity as the sole predictor, without adjustment for covariates. The overall pattern of findings remained largely unchanged, although several additional associations reached significance in the unadjusted models (Tables S1 and S2).

### 2.5. Pre-processing

Data preprocessing and statistical analyses were conducted in MATLAB (Kempler et al., 2024), with statistical significance set at α = .05. Bivariate Pearson correlations were first computed across all study measures to characterize zero-order relationships among misophonia severity, ToM and empathy performance along with reflective functioning, alexithymia, mimicry, sound sensitivity, and anxiety/depressive symptoms. All task and questionnaire responses were complete because forced-response settings were used. Age and/or sex information was unavailable for 12 participants; therefore, covariate-adjusted analyses including these demographic variables were conducted on 329 participants.

For behavioural outcomes, accuracy and RT indices were derived from the Yoni and MET tasks after excluding practice and attention-check trials. The performance was indexed by the proportion of correctly identified responses and by RT for correct trials as a secondary index of processing efficiency. In the Yoni task, accuracy and RT were calculated overall and separately by mentalizing order (first- vs second-order) and condition (cognitive vs affective ToM). In the MET, accuracy and RT were calculated overall and separately for positive- and negative-valence stimuli (Dziobek et al., 2008; Isernia et al., 2023). Extreme RT values were screened at the trial level, and observations shorter than <200lllms were excluded. Questionnaire scores were retained unless there was evidence of invalid responding, as extreme values were considered potentially meaningful expressions of the underlying phenotype rather than measurement error. No participants met the aforementioned criteria. Participants were additionally classified as meeting the established S-Five cutoff for clinically elevated misophonia at scores ≥87 (Vitoratou et al., 2023), although the primary analyses treated misophonia severity dimensionally.

We also conducted sensitivity analyses to assess the influence of potentially influential observations on the primary S-Five models. For multiple linear regression (MLM), cases were flagged using |studentized residual| > 3 or Cook’s D > 4/N; analogous influence diagnostics were applied to logistic models. Models were then refitted after excluding flagged cases, and robustness was evaluated by comparing coefficient direction and false discovery rate (FDR)-adjusted significance with the full-sample results (Bollen et al., 1985; Cook, 2000).

### 2.6. Model Specification (Multiple Linear Modelling)

For inferential analyses, MLM modelling approaches were used to characterize the social-cognitive attributes of misophonia severity. MLMs examined associations between misophonia severity (S-Five total score and its five factors) and individual-level indices of mentalizing (Yoni task), empathy (MET and QCAE), reflective functioning (RFQ-8), alexithymia (TAS-20), and mimicry (IMQ). All models adjusted for age, sex, sound sensitivity (SSSQ), and symptoms of anxiety and depression (SAD-T), thereby testing whether associations with misophonia severity were evident beyond these potentially overlapping sources of individual variation.

For the Yoni and MET, linear mixed-effects models (LMERs) were additionally used to a) account for within-participant variability across task conditions and to b) characterize both the main effects of misophonia severity and its interactions with theoretically relevant within-task dimensions. Specifically, separate models examined interactions between S-Five total scores and Yoni mentalizing order (first- and second-order), Yoni condition (cognitive ToM, affective ToM, and physical control), and MET stimulus valence (positive and negative) for both accuracy and RT. Participant-level random intercepts accounted for the non-independence of repeated observations within individuals. In addition, a binary logistic regression model examined whether misophonia severity (S-Five total score) predicted the likelihood of endorsing mimicry (IMQ: no = 0, yes = 1), adjusting for age, sex, sound sensitivity, and symptoms of anxiety and depression.

Standardized regression coefficients (β) and p values are reported throughout. To limit inflation of Type I error across related analyses, p values were additionally adjusted using the Benjamini–Hochberg FDR procedure (Benjamini et al., 1995) and was reported across all analysis.

## 3. Results

### 3.1. Correlation Between Study Variables

Zero-order correlations revealed a coherent but differentiated pattern across behavioural and self-report measures. Greater S-Five severity was correlated with poorer Yoni and MET performance, reduced reflective-functioning certainty, greater reflective-functioning uncertainty, higher alexithymia, and elevated sound-sensitivity and anxiety/depressive symptoms. Behavioural accuracy and RT indices were also intercorrelated across tasks, whereas several questionnaire domains showed only weak or nonsignificant associations (Figure 2A).

**Figure 2.**
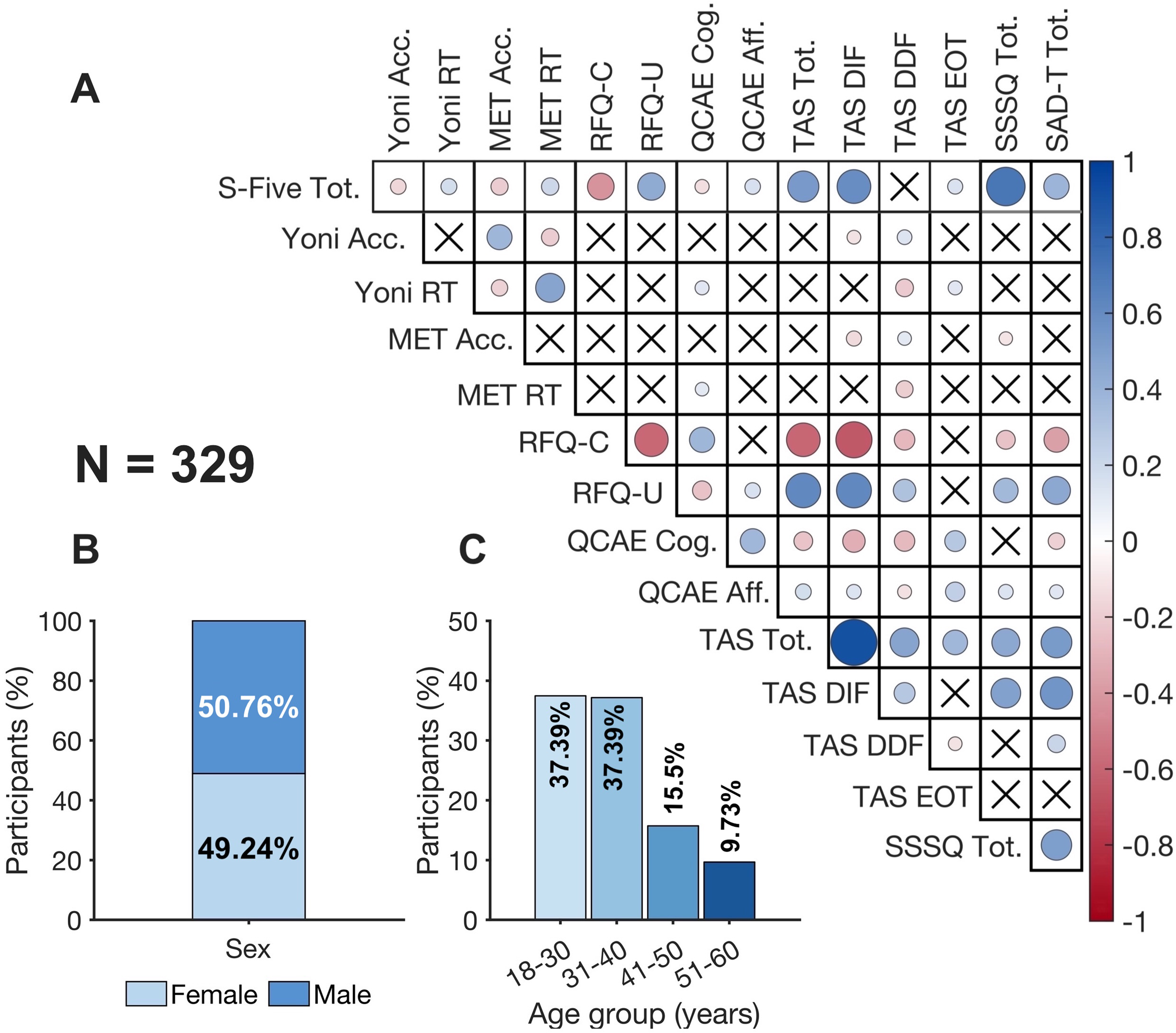
Sample composition and zero-order associations. Pearson correlations among misophonia severity and other study variables (N = 341). Circle size and saturation indicate correlation magnitude; blue denotes positive and red negative associations. Crosses indicate p ≥ .05. (B) Sex distribution. (C) Age-group distribution (N = 329). Acc., accuracy; RT, response time; RFQ-C/U, certainty/uncertainty; QCAE Cog. /Aff., cognitive/affective; TAS DIF/DDF/EOT, difficulty identifying feelings/difficulty describing feelings/externally oriented thinking; SSSQ, sound sensitivity symptoms Questionnaire; SAD-T, Screening for anxiety and depressive symptoms in tinnitus-hyperacusis-misophonia.

The sex distribution was approximately balanced, with 49.24% female and 50.76% male participants (Figure 2B). The age distribution was skewed toward younger participants, with comparatively fewer individuals in the older age groups (Figure 2C).

### 3.2. S-Five is associated with decreased performance on Yoni and MET tasks, increased TAS and altered RFQ

Higher S-Five Total scores were associated with a broad but selective pattern of social-cognitive differences after adjustment for age, sex, sound sensitivity, and anxiety and depressive symptoms (Figure 3A). In more details, increased misophonia severity appeared to be associated with lower Yoni total accuracy, β = −.217, p = .006, pFDR = .017, and slower Yoni RT, β = .194, p = .012, pFDR = .028. A similar pattern was observed for the MET, with higher S-Five scores associated with lower total accuracy, β = −.205, p = .006, pFDR = .017, and slower total RT, β = .327, p < .001, pFDR < .001 (Figure 3B–3E). For the distribution of accuracy and RTs of Yoni and MET as well as analyses examining the contributions of the individual S-Five factors, see Figures S1 and S2, S3, S6 and S7.

**Figure 3.**
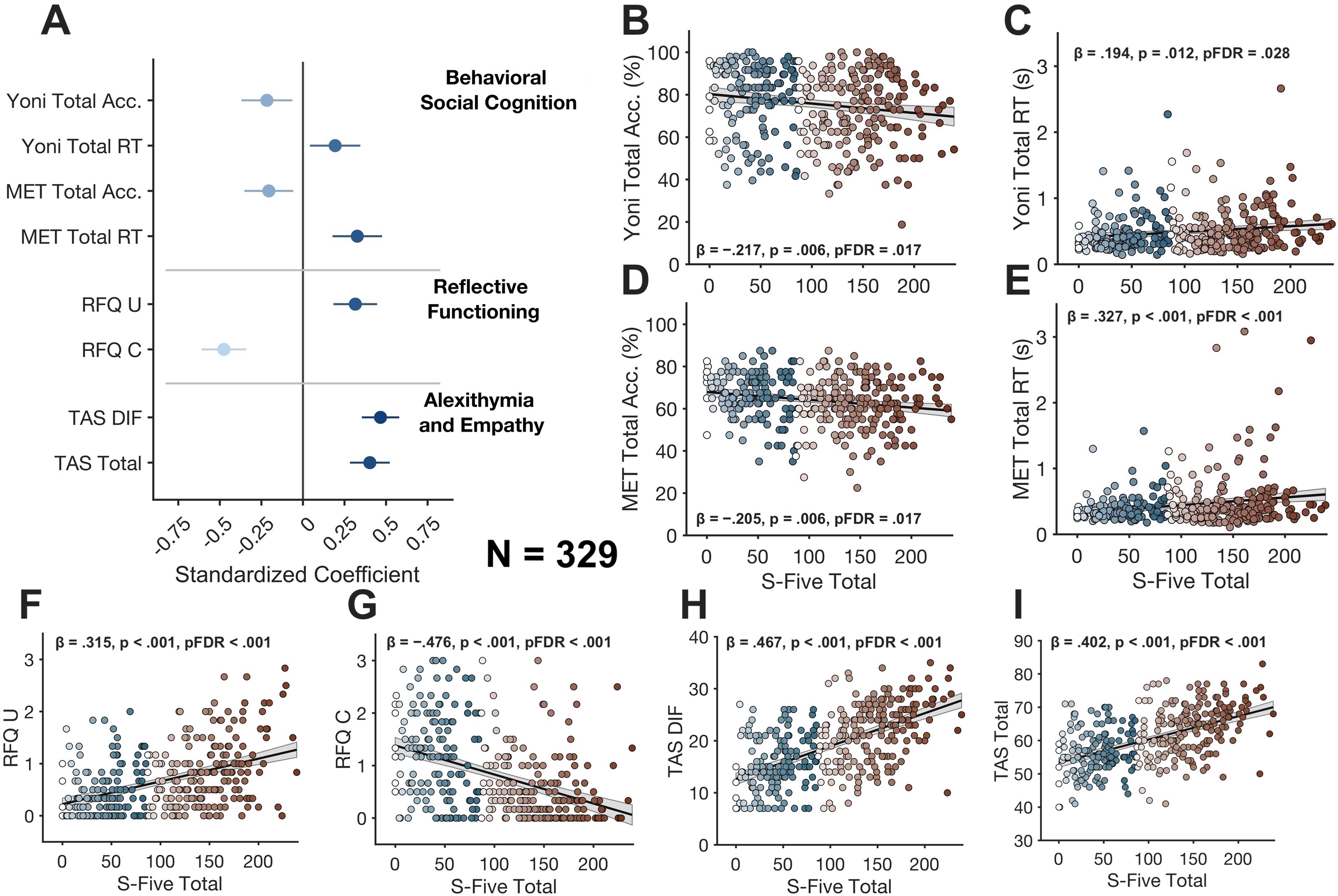
Associations between S-Five Total scores and social-cognitive, reflective-functioning, alexithymia, and empathy outcomes (N = 329). (A) Adjusted standardized regression coefficients for associations between S-Five Total and Yoni and MET performance, reflective functioning (RFQ), and alexithymia (TAS-20), controlling for age, sex SSSQ, and SAD-T. Horizontal lines indicate 95% confidence intervals. (B–E) Descriptive, unadjusted associations between S-Five total and Yoni total accuracy and RT and MET total accuracy and RT. (F–I) Descriptive, unadjusted associations with RFQ-Certainty (RFQ-C), RFQ-Uncertainty (RFQ-U), TAS Difficulty Identifying Feelings (TAS DIF), and TAS Total, respectively. Black lines show unadjusted linear fits and shaded areas indicate 95% confidence intervals. Points below the S-Five cutoff of 87 are shown in blue and those at or above 87 in orange to red, with darker shades indicating higher S-Five scores within each range. Bold annotations report adjusted standardized coefficients (β) and Benjamini–Hochberg FDR-adjusted p values. RTs are shown in seconds (N = 329).

The association with Yoni accuracy was concentrated in second-order, affective, and cognitive mentalizing. Higher S-Five Total scores were associated with lower second-order accuracy, β = −.24, p = .002, pFDR = .008; lower affective ToM accuracy, β = −.22, p = .005, pFDR = .017; and lower cognitive ToM accuracy, β = −.214, p = .006, pFDR = .017. For RT, higher severity was only associated with slower affective ToM performance, β = .194, p = .013, pFDR = .028 (See Figures S2, S3, S4 and S5 for all models across constructs) .

For the MET, valence-specific analyses showed lower accuracy for only positive stimuli, β = −.246, p < .001, pFDR = .004. However, higher severity was related to slower RTs for both positive stimuli, β = .304, p < .001, pFDR < .001, and negative stimuli, β = .258, p < .001, pFDR = .004 (also see Figures S6 and S7).

In addition, the mixed-effects analyses provided little evidence that the association between misophonia severity and task performance differed across within-task dimensions. None of the S-Five-by-task-factor interactions remained significant after FDR correction. For the Yoni task, the S-Five-by-order interaction for accuracy approached significance after correction, F(1, 329) = 6.00, p = .015, pFDR = .059, whereas the interaction for RT was not significant, F(1, 329) = 0.24, p = .627, pFDR = .627. S-Five-by-condition interactions were also nonsignificant for Yoni accuracy, F(2, 658) = 0.88, p = .417, pFDR = .556, and RT, F(2, 651.39) = 1.75, p = .175, pFDR = .351. Similarly, S-Five-by-valence interactions were nonsignificant for MET accuracy, F(1, 329) = 0.95, p = .331, pFDR = .331, and RT, F(1, 329) = 1.12, p = .291, pFDR = .331.

Moreover, greater misophonia severity was also associated with poorer self-reported mentalizing. S-Five Total scores were positively associated with uncertainty about mental states, β = .315, p < .001, pFDR < .001, and negatively associated with certainty about mental states, β = −.476, p < .001, pFDR < .001. Higher severity was further associated with greater DIF, β = .467, p < .001, pFDR < .001, and higher total alexithymia, β = .402, p < .001, pFDR < .001 (Figure 3F-3I) (Figure S8-S11).

Overall, the findings indicate a selective profile characterized by reduced behavioural efficiency, altered reflective functioning, heightened alexithymia, and increased mimicry, rather than a uniform impairment across all domains of social cognition.

### 3.3. S-Five is Linked to Higher Probability of Mimicry Endorsement

We observed a relationship between higher S-Five total score and a greater likelihood of endorsing mimicry of trigger-producing actions or sounds, standardized log-odds coefficient = .319, p < .001, pFDR < .001, after adjustment for age, sex, sound sensitivity, and anxiety and depressive symptoms (Figure 4A). Mimicry data were available for 332 participants, 89 participants (26.8%) endorsed mimicry, whereas 243 (73.2%) did not (Figure 4B). Among participants with clinically elevated S-Five scores (≥87; N = 188), 77 (41.0%) endorsed mimicry and 111 (59.0%) did not (Figure 4C).

**Figure 4.**
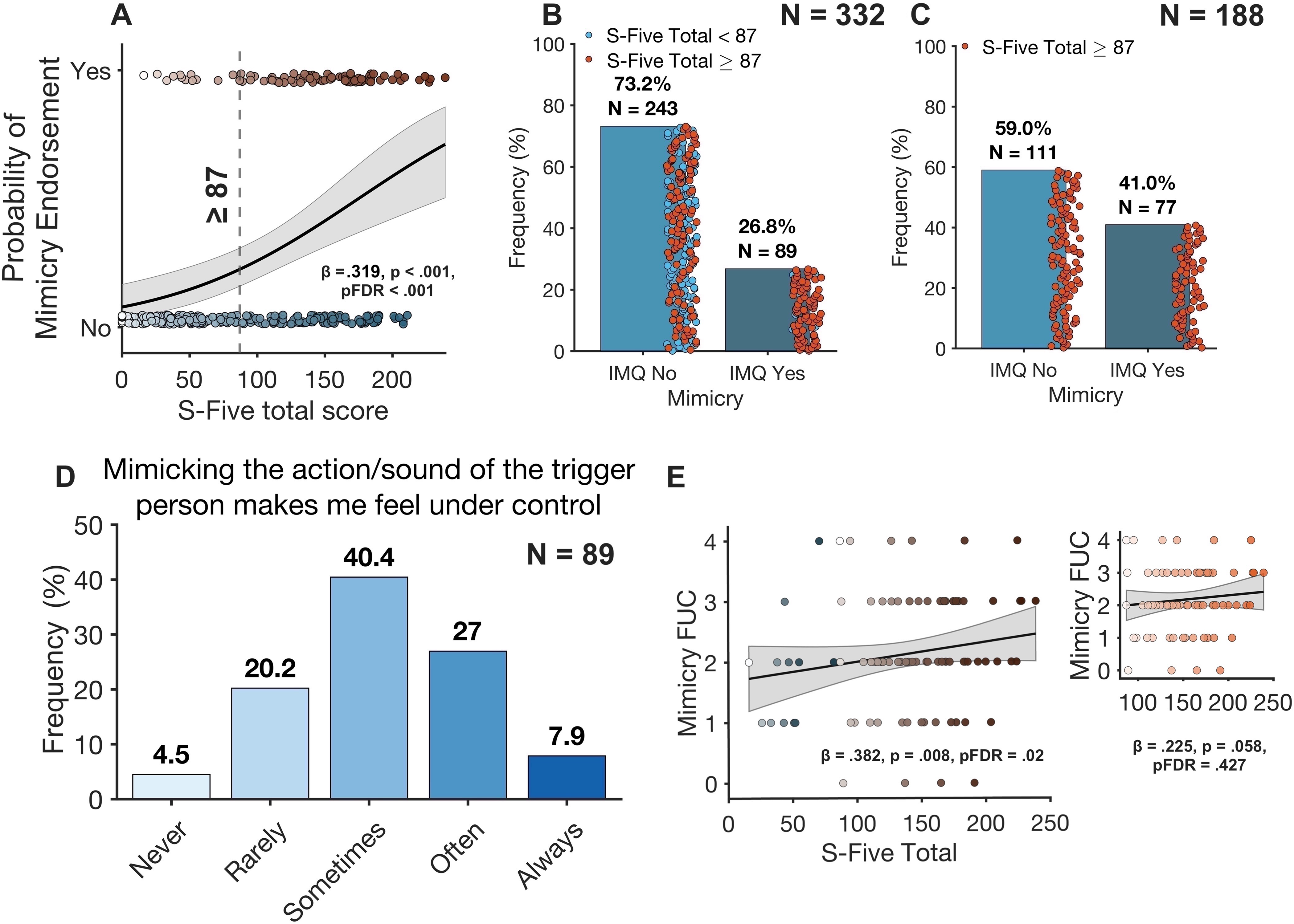
Misophonia severity and mimicry. Predicted probability of mimicry endorsement across S-Five Total scores; shading indicates the 95% confidence interval, and the dashed line marks the ≥87 cutoff. (B) Mimicry endorsement in the sample (N = 332). (C) Mimicry endorsement among participants meeting the S-Five ≥87 cutoff (N = 188). (D) Reported frequency with which mimicry increased perceived control among endorsers (N = 89). (E) Association between S-Five Total scores and reported frequency of mimicry as a means of gaining control; the line denotes the fitted regression with 95% confidence interval. The inset depicts the same association only in individuals with S-Five ≥87 cutoff.

Among all participants endorsing mimicry, greater S-Five total scores were linked to more frequent reports that mimicry restored a sense of control, β = .382, p = .008, pFDR = .02 (Figure 4E). The associations with relief from distress, β = .315, p = .025, pFDR = .052, spontaneous mimicry, β = .175, p = .195, pFDR = .252, and deliberate mimicry, β = −.031, p = .827, pFDR = .850, were not significant. Thus, mimicry was more prevalent at higher levels of misophonia severity and was associated with a perceived restoration of control.

Among participants who endorsed mimicry (N = 89), 4.5% reported they feel more in control, 20.2% rarely, 40.4% sometimes, 27.0% often, and 7.9% always (Figure 4D). For other mimicry components see Figures S12A- S12C.

The primary analyses and objectives of the present study focused on overall misophonia severity, indexed by the S-Five Total score. Analyses examining associations across the individual S-Five factors are reported in the supplementary materials.

### 3.4. Sensitivity Analysis

The principal findings were largely unchanged after exclusion of influential observations. Associations involving Yoni total accuracy and RT, MET total accuracy and RT, RFQ-certainty (RFQ-C) and -uncertainty (RFQ-U), TAS DIF, total alexithymia, and mimicry endorsement remained significant. Overall, these findings were robust to the exclusion of influential observations. Full sensitivity results are reported in Table S3.

## 4. Discussion

This study provides convergent evidence that misophonia severity is associated with a selective alteration in social-cognitive functioning rather than an only auditory deficit. Heightened S-Five severity was linked to lower accuracy and slower responding on the Yoni task and MET, indicating reduced efficiency in mental-state inference and empathic functioning. At the self-report level, higher severity was associated with lower certainty and greater uncertainty about mental states, increased difficulty identifying feelings, and higher overall alexithymia. On the other hand, global cognitive and affective empathy were largely preserved. Misophonia severity was also associated with a greater likelihood of mimicking trigger-producing actions or sounds, particularly when mimicry was experienced as restoring a sense of control.

These findings corroborate a model in which misophonia involves altered self–other processing across partially distinct levels, interpretation of others’ mental and emotional states, representation of one own internal state, and embodied regulatory responses to socially generated triggers. These associations remained after accounting for age, sex, anxiety, sound sensitivity and depressive symptoms, suggesting that they are not readily reducible to general distress or abnormal sound tolerance. However, as the data are cross-sectional, the findings identify an interrelated social-cognitive profile rather than establishing causal mechanisms which will be discussed in detailed with its caveats.

### 4.1. Misophonia Severity Tracks Reduced ToM and Mentalizing

Misophonia severity was associated with lower accuracy across both cognitive and affective ToM, alongside slower overall responding, indicating less efficient mental-state inference. This extends evidence that misophonic reactions are shaped by the social meaning of triggers, including the identity and perceived intentions of the sound source (Berger et al., 2024; Humolli et al., 2025; Siepsiak et al., 2023).

At the order level, higher misophonia severity was related to lower second-order, but not first-order, Yoni accuracy. This suggests that basic inference about another person’s immediate thoughts or intentions may remain relatively intact (i.e., “He is chewing because he is hungry.”), whereas more recursive judgments, representing what one person thinks about another person’s mental state, may be more vulnerable (i.e., “He knows that I find chewing distressing, but he thinks I am overreacting, so he does not expect me to be upset.”). However, this apparent second-order selectivity should be interpreted cautiously, as the S-Five-by-order interaction was not significant and therefore did not confirm a stronger severity effect for second- than first-order mentalizing. This pattern fits well with hierarchical models of mentalizing in which second-order ToM places greater demands on coordinating nested mental representations (Isernia et al., 2023; Schurz et al., 2021). At the neural level, higher-order mentalizing and misophonia implicate partially overlapping medial prefrontal–insular circuitry, suggesting potential neurofunctional convergence between higher-order social inference and misophonic processing (Kumar et al., 2021; Kumar et al., 2017; Neacsiu et al., 2026; Zhen et al., 2021).

These findings were further supported by lower reflective-functioning certainty and higher uncertainty about mental states. This may be particularly salient in misophonia, where trigger aversiveness is modulated by the social attribution and inferred intentionality of the sound source (Humolli et al., 2025; Siepsiak et al., 2023). Uncertainty in mentalizing may therefore make the intentions underlying trigger-producing behaviour harder to represent, increasing its social ambiguity and salience.

Our observations point to the view that misophonia is accompanied by differences in the representation and interpretation of mental states, rather than by a task-specific decrement alone (Fonagy et al., 2016). This interpretation complements neurobiological evidence that misophonia recruits distributed salience, motor, and social-perceptual systems beyond auditory pathways (Kumar et al., 2021; Kumar et al., 2017). As the present measures were administered outside a trigger context, however, they are best viewed as indexing a more general social-cognitive phenotype of misophonia rather than the proximal processes that generate trigger reactivity (Black et al., 2025).

### 4.2. Greater Misophonia Severity Characterize Abnormality in Empathic Processing and Elevated Alexithymia

The findings showed a relationship between misophonia severity and lower empathy-task accuracy and prolonged response latencies, pointing to a level of impairment in decoding of emotional states. Accuracy effects were evident for positive, but not negative, stimuli, whereas response slowing generalized across both valence categories. However, the absence of a significant S-Five-by-valence interaction indicates that these effects did not differ statistically by emotional valence. This accords with general notion of altered emotional processing related to misophonia (Black et al., 2025; Hanna et al., 2026; Kumar et al., 2017; Scheerer et al., 2026), but contrasts with Koroglu Gokbel et al. (2026). They reported that increased self-reported emotional empathy, indexed by the Cognitive and Affective Empathy Scale, was related to higher misophonia severity. The divergence may reflect the distinction between behavioural proficiency in emotion decoding and subjectively reported empathic responsivity, which need not vary in parallel (Black et al., 2025).

Poorer empathy-related performance in misophonia arises not from attenuated emotional responsivity, but from excessive weighting of self-relevant affective and interoceptive signals. The anterior insula integrates bodily arousal, affective salience, and socially relevant emotional information, and is prominently recruited during empathic processing (Gu et al., 2012; Lamm et al., 2010). In misophonia, heightened anterior-insula responsivity may therefore amplify the listener own aversive state, potentially drawing processing resources toward self-oriented distress rather than precise representation of another person’s emotion. Under conditions of heightened arousal, these competing demands on limited processing resources may further constrain other-oriented appraisal. Empathy depends not only on affective sharing but also on maintaining self–other distinction and regulating one’s own response; when self-focused arousal dominates, empathic concern can give way to personal distress and less effective other-oriented processing (Decety et al., 2006; Lamm et al., 2007).

The elevated alexithymia observed here lends further support to this finding. Difficulty identifying one own affective state can coexist with strong physiological reactivity and has been linked to poorer empathy and altered insular responses during observation of others’ emotions (Bird et al., 2010; Moriguchi et al., 2007). Moriguchi et al. (2007) reported enhanced right-insula activation alongside lower empathy-related scores in alexithymic individuals, illustrating that greater insular recruitment does not necessarily translate into more effective empathic understanding. This is particularly pertinent to misophonia, where trigger exposure has been shown to elicit exaggerated anterior-insula responsivity (Kumar et al., 2017; Schroder et al., 2019).

### 4.3. Mimicry may Function as an Embodied Regulatory Response in Misophonia

Higher likelihood of mimicking trigger-producing actions or sounds was associated with the severity of misophonia, and among endorsers, more frequent reports that mimicry restored a sense of control. Out of 188 participants meeting the ≥ 87 cutoff score established by Vitoratou et al. (2023), 41% endorsed mimicry. This build on prior evidence that mimicry is common in misophonia and covaries with symptom severity and trigger type (Ash et al., 2023; Edelstein et al., 2013; Rouw et al., 2018).

One interpretation that was put forward by Kumar et al. (2021), is that mimicry in misophonia may derive from heightened recruitment of motor representations corresponding to trigger-producing actions. Since many common triggers involve human orofacial behaviours such as chewing and eating, these sounds may automatically recruit the somatotopically corresponding orofacial region of ventral premotor cortex, which Kumar et al. (2021) found to be selectively hyperactive during trigger exposure and more strongly coupled with auditory cortex. This neural profile was interpreted as “hyper-mirroring,” whereby the actions of others are excessively represented within the observer’s own motor system, with sound serving as the medium through which those actions are mirrored. Rather than reflecting an exaggerated auditory response per se, the trigger may therefore evoke an unusually strong internal motor representation of the action that produced it. This excessive motor representation may, in turn, promote spontaneous mirroring of the trigger-producing action (Kumar et al., 2021).

A second and mutually inclusive possibility was proposed by Ash et al. (2023) suggesting that overt mimicry may function as a regulatory response to an excessive spontaneous urge to mirror the trigger-producing action. By deliberately reproducing the same action, the individual may generate a more predictable sensory consequence and regain a sense of agency over an otherwise externally driven event. In this framework, greater predictability and control may reduce the mismatch between expected and incoming sensory input, thereby attenuating distress (Ash et al., 2023). Echoing this pattern, exaggerated perceived control in objectively uncontrollable social interactions and heightened aversion to unexpected social events has been reported in misophonia (Banker et al., 2022).

A third possibility is that heightened mimicry indexes greater reliance on embodied social processing. Perception-action models posit that observing another person behaviour automatically evokes corresponding motor representations, providing a low-level route through which others’ actions and affective states can be represented (Prochazkova et al., 2017; Rizzolatti et al., 2004). In the present study context, increased mimicry alongside less efficient ToM, empathy and emotion decoding may as the result indicate a relative shift toward embodied resonance when explicit social-cognitive inference is less efficient which we consider it as “compensatory mimicry”. Berger et al. (2024) similarly proposed “motor empathy” as a potentially relevant dimension of misophonia.

### 4.4. Integrated Conceptual Model

Taken everything into account, the findings substantiate a social-cognitive dysregulation model in which misophonia is associated with altered processing across multiple levels of self–other representation. Increased symptom severity was linked to reduced clarity about less efficient decoding of others’ intentions and emotions, greater uncertainty in representing mental states, one’s own affective states, and increased mimicry of trigger-producing actions. These components are conceptually different but potentially interacting, imprecise self-state representation may heighten ambiguity during social appraisal, while greater processing demands during mental-state inference may increase the salience of socially generated triggers (Bird et al., 2014; Fonagy et al., 2016; Schurz et al., 2021). Mimicry may then operate as an embodied attempt to restore predictability or control. This model extends auditory-affective accounts of misophonia by incorporating reflective, inferential, and sensorimotor processes, without implying that any single component is necessary or sufficient.

### 4.5. Limitations

Several limitations constrain interpretation. First, the cross-sectional design precludes causal inference and cannot determine whether social-cognitive differences precede, maintain, or result from misophonic symptoms. Second, several constructs were assessed by self-report and may be affected by shared-method variance and response bias. Third, online administration reduced experimental control over testing conditions, although it enabled a large sample and standardized task delivery. Fourth, the large number of outcomes necessitated multiple-testing correction using the Benjamini–Hochberg false discovery rate procedure, which may have reduced sensitivity to weaker associations. Fifth, clinical attributes were based primarily on self-report rather than structured diagnostic assessment. Finally, the Yoni and MET tasks were completed outside a trigger context; therefore, the study cannot establish whether the observed social-cognitive inefficiencies are amplified during misophonic arousal. At the same time, their presence in the absence of explicit triggers suggests that misophonia-related differences may extend beyond immediate sound-evoked processing to broader aspects of social and emotional functioning.

### 4.6. Conclusion

The present findings suggest that misophonia may involve selective alterations in self–other processing in addition to abnormal auditory-affective reactivity. The most consistent associations concerned behavioural social-cognitive efficiency, emotional self-identification, reflective functioning, and mimicry, whereas global empathy was largely preserved. This profile argues against a unitary social-cognitive deficit and instead supports a multidimensional account in which auditory, affective, cognitive mentalizing, and embodied regulatory processes interact. For future research, it would also be fruitful to investigate the physiological mechanisms underlying the observed deficits, as this could help refine and strengthen the proposed model. In addition, experimental studies are needed to determine whether trigger exposure magnifies these effects and to evaluate whether social-cognitive processes represent mechanisms, consequences, or compensatory responses within misophonia.

## Acknowledgement

We would like to thank SoQuiet and Duke Centre for Misophonia and Emotion Regulation (CMER). This study has been funded by ESSCA School of Management.

## CRediT author statement

Mercede Erfanian: Conceptualization, Data Curation, Formal Analysis, Funding Acquisition, Investigation, Methodology, Project Administration, Resources, Software, Supervision, Validation, Visualization, Writing – Original Draft Preparation, Writing – Review & Editing. Sukhbinder Kumar: Conceptualization, Resources, Supervision, Writing – Review & Editing.

---

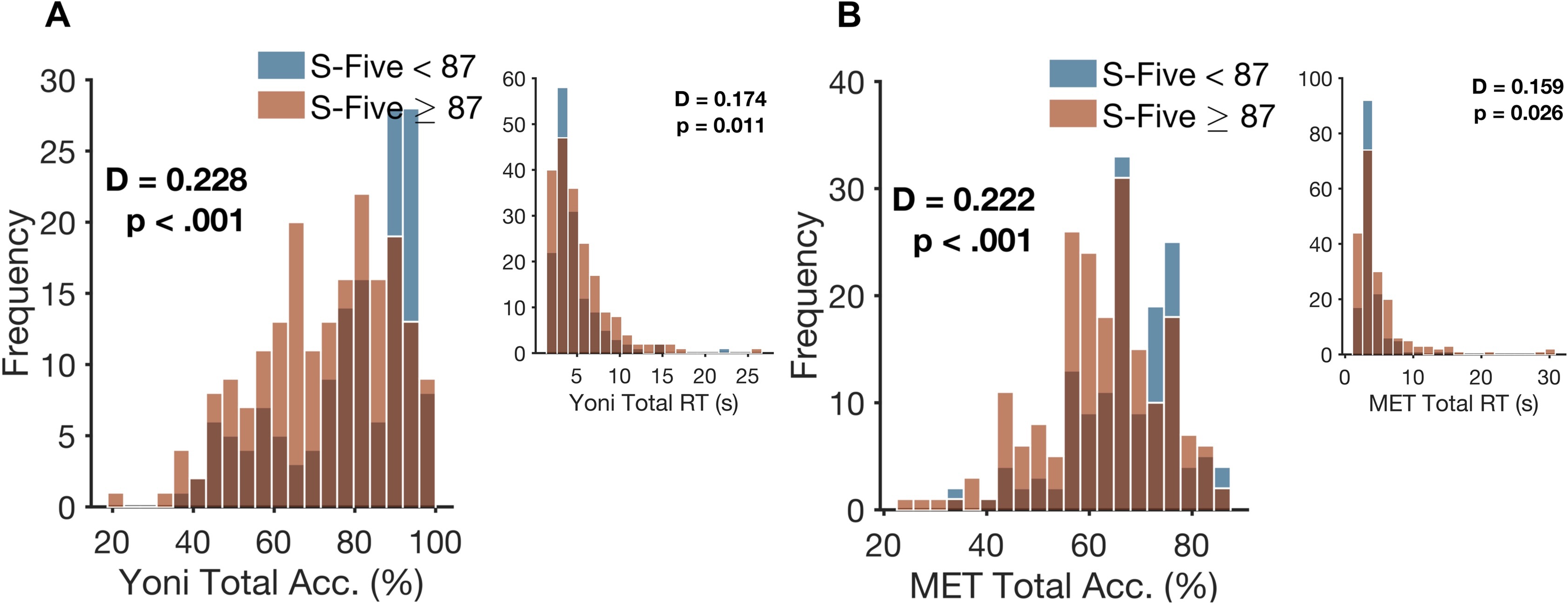

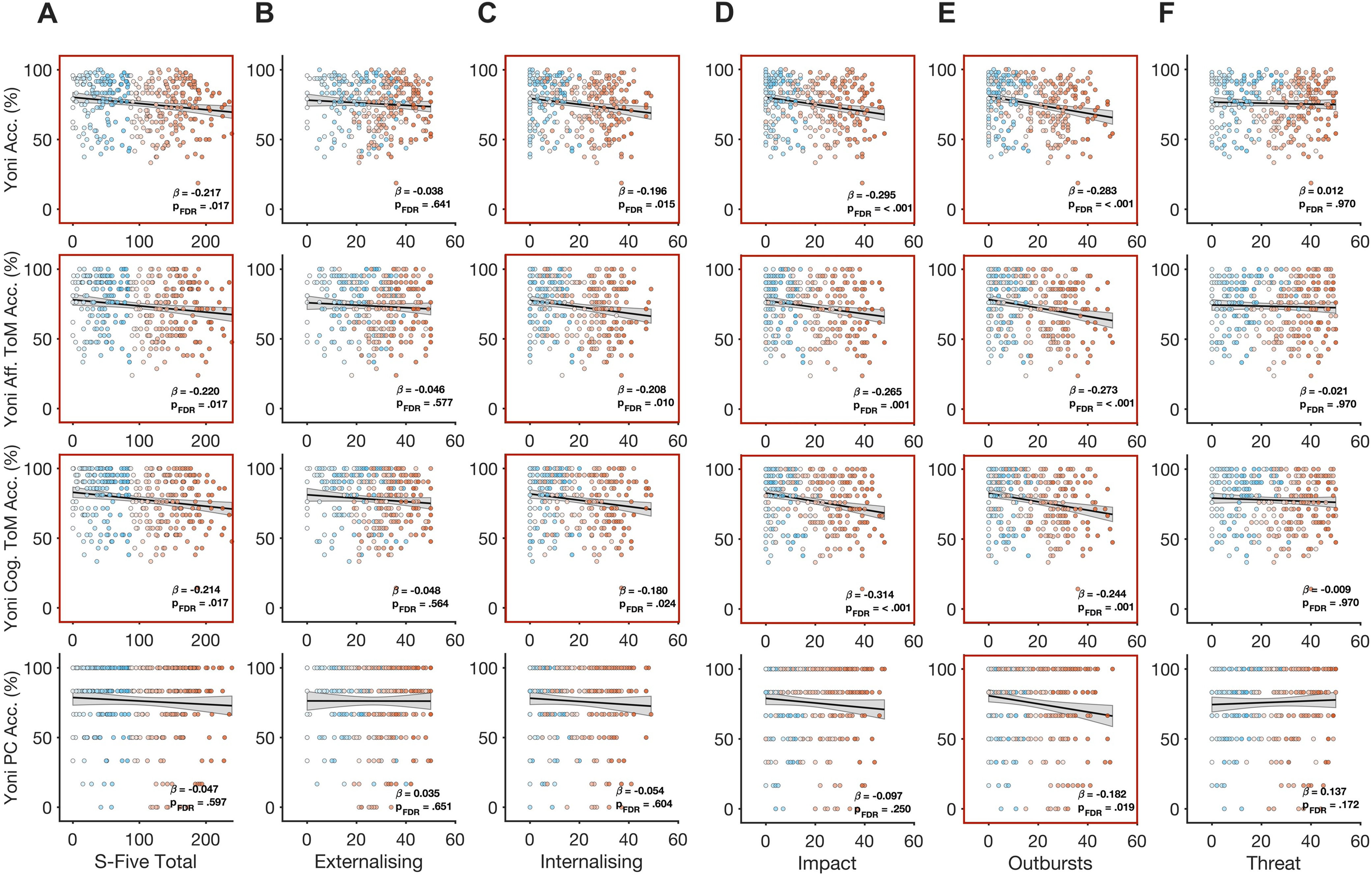

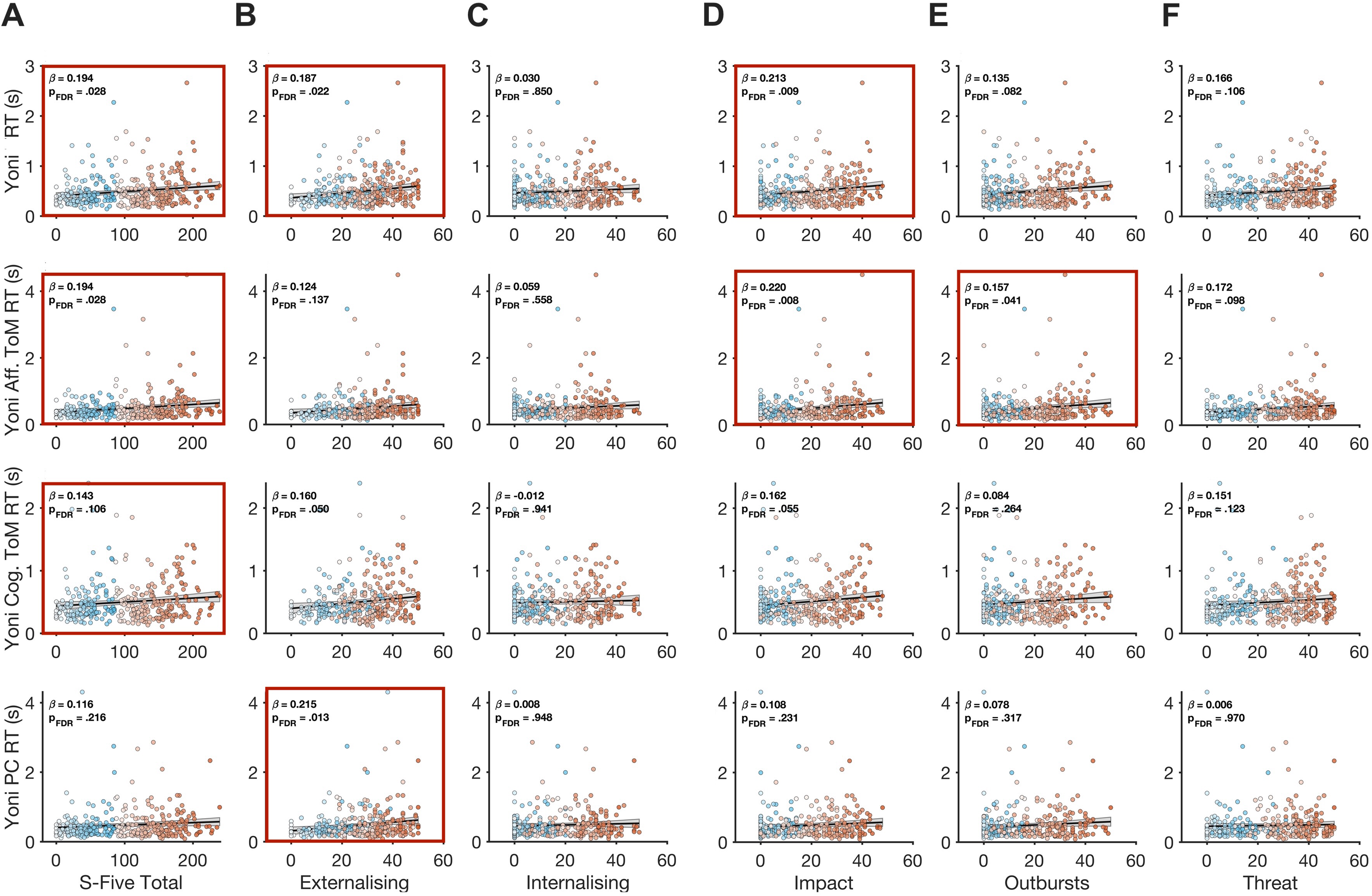

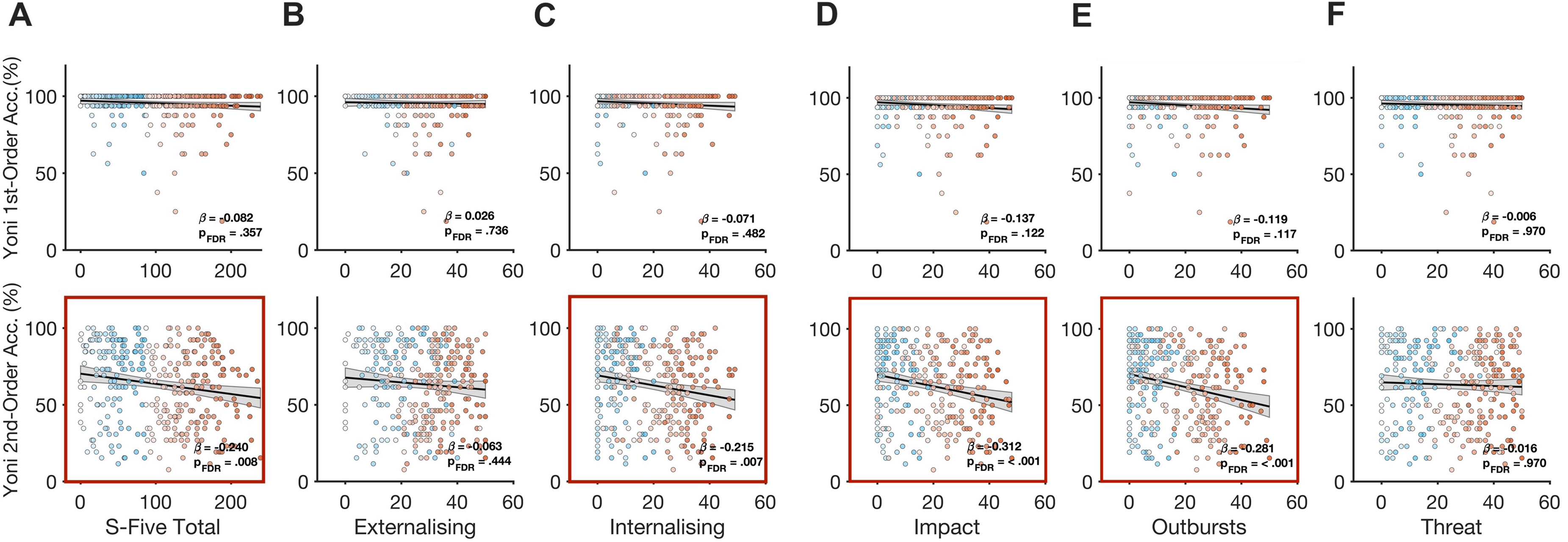

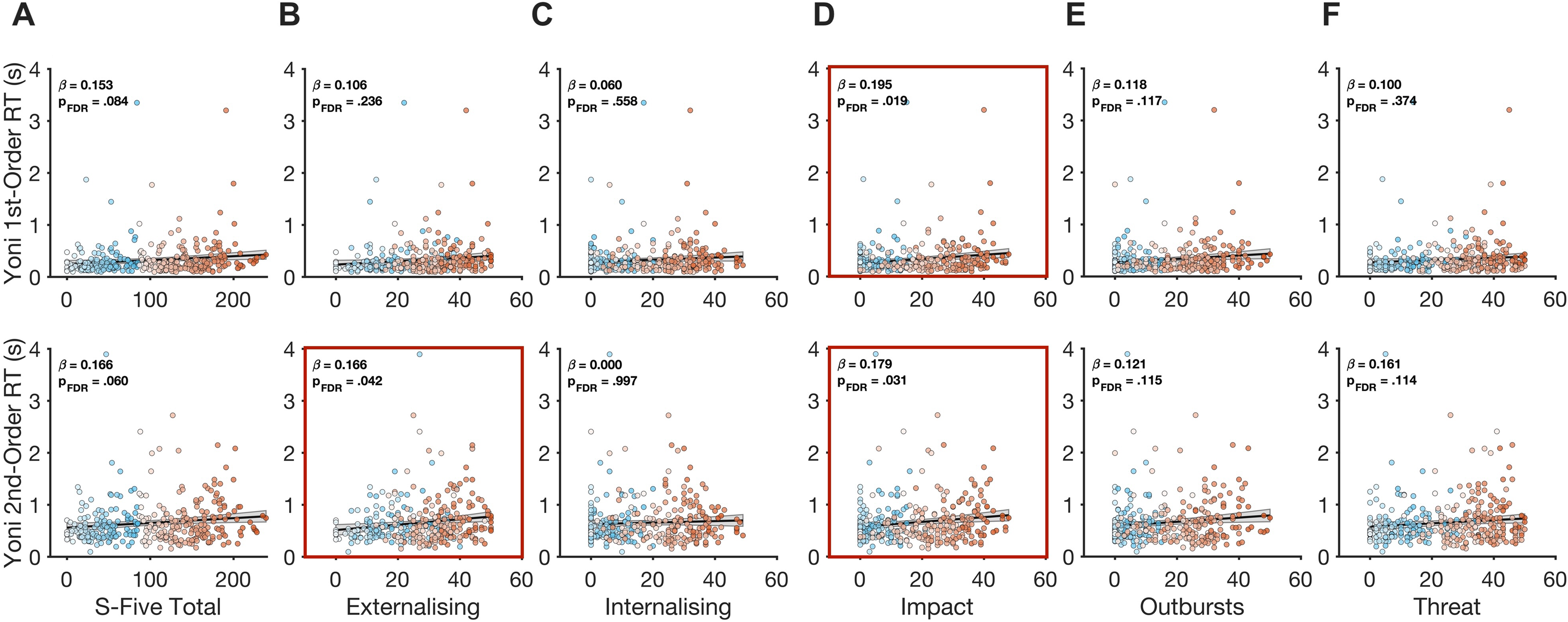

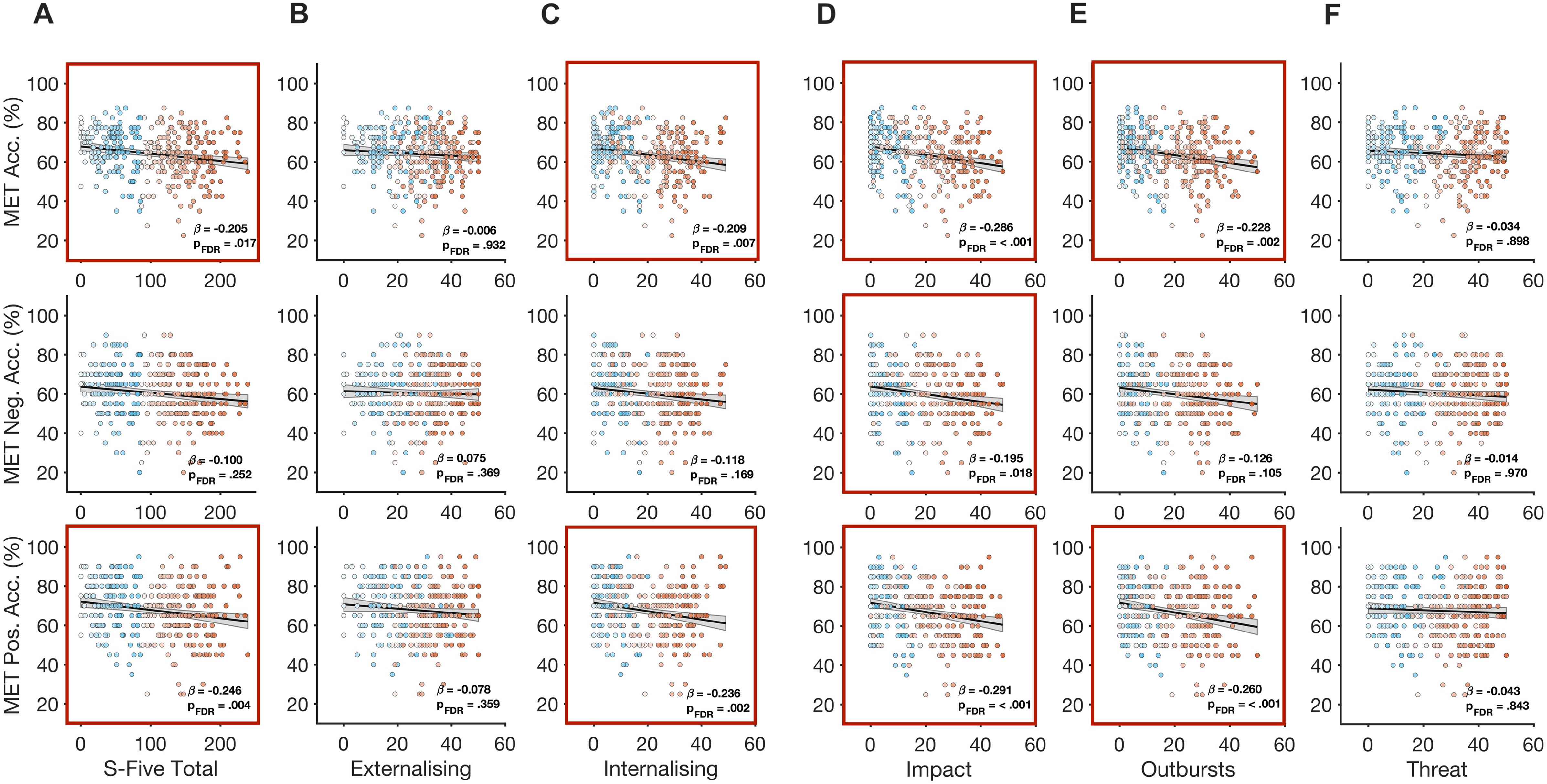

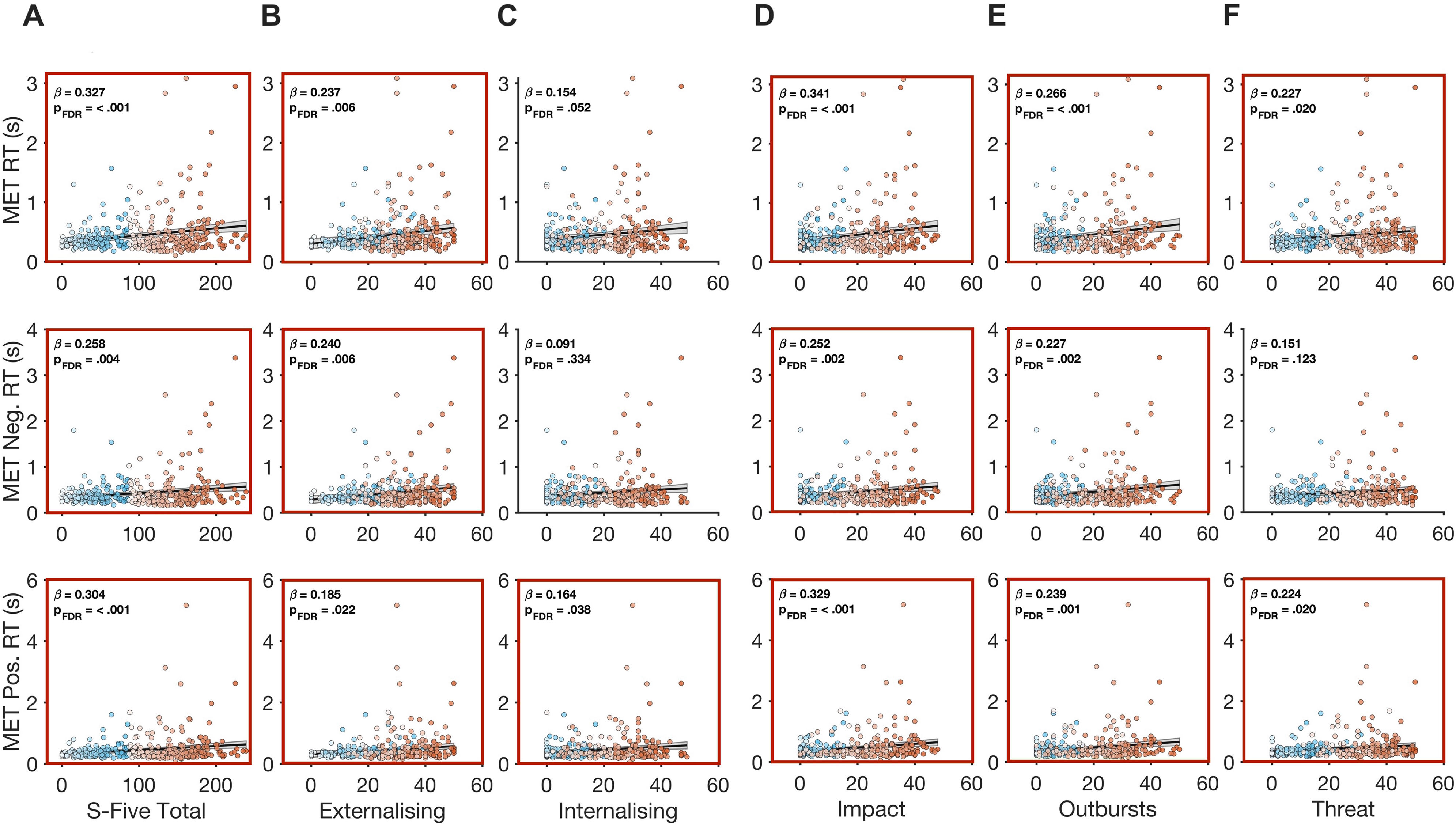

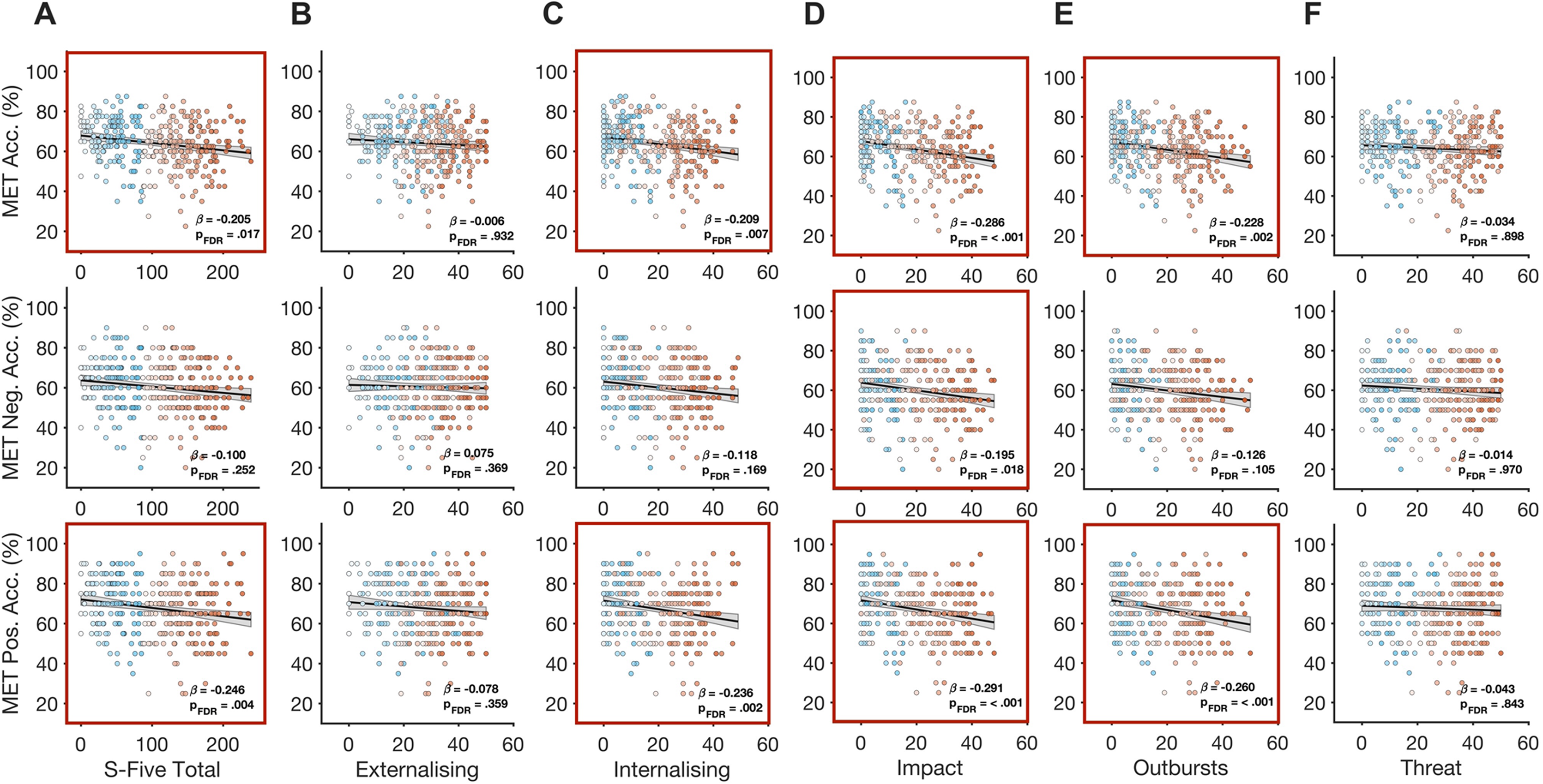

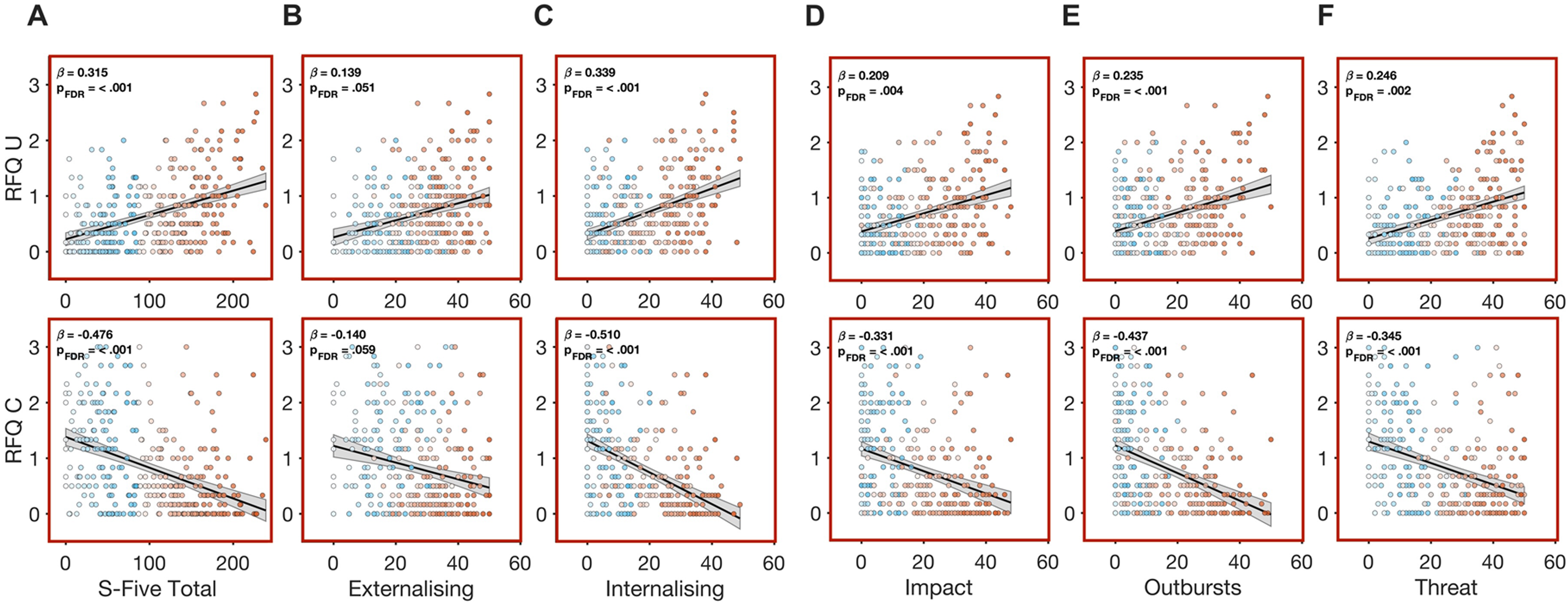

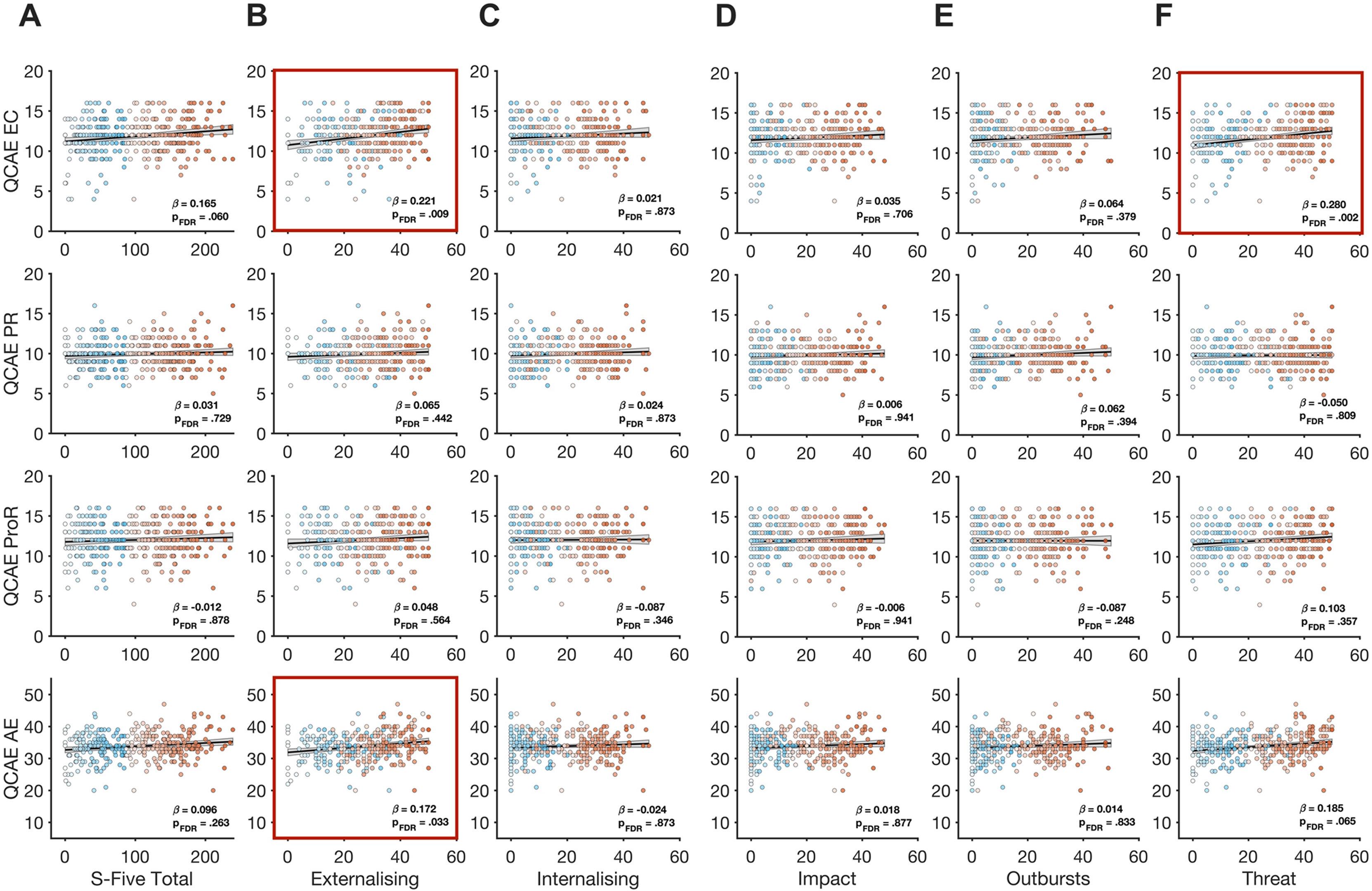

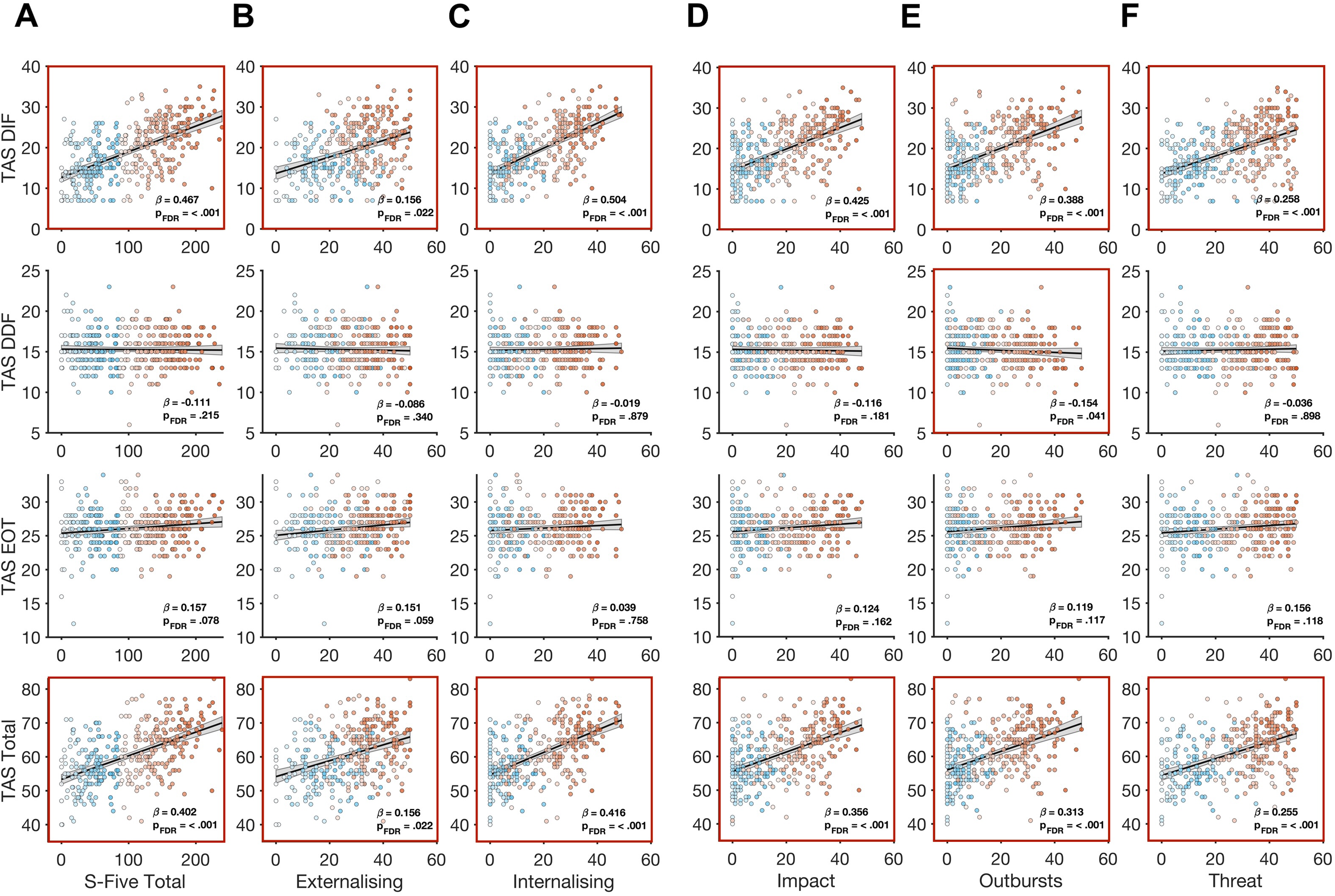

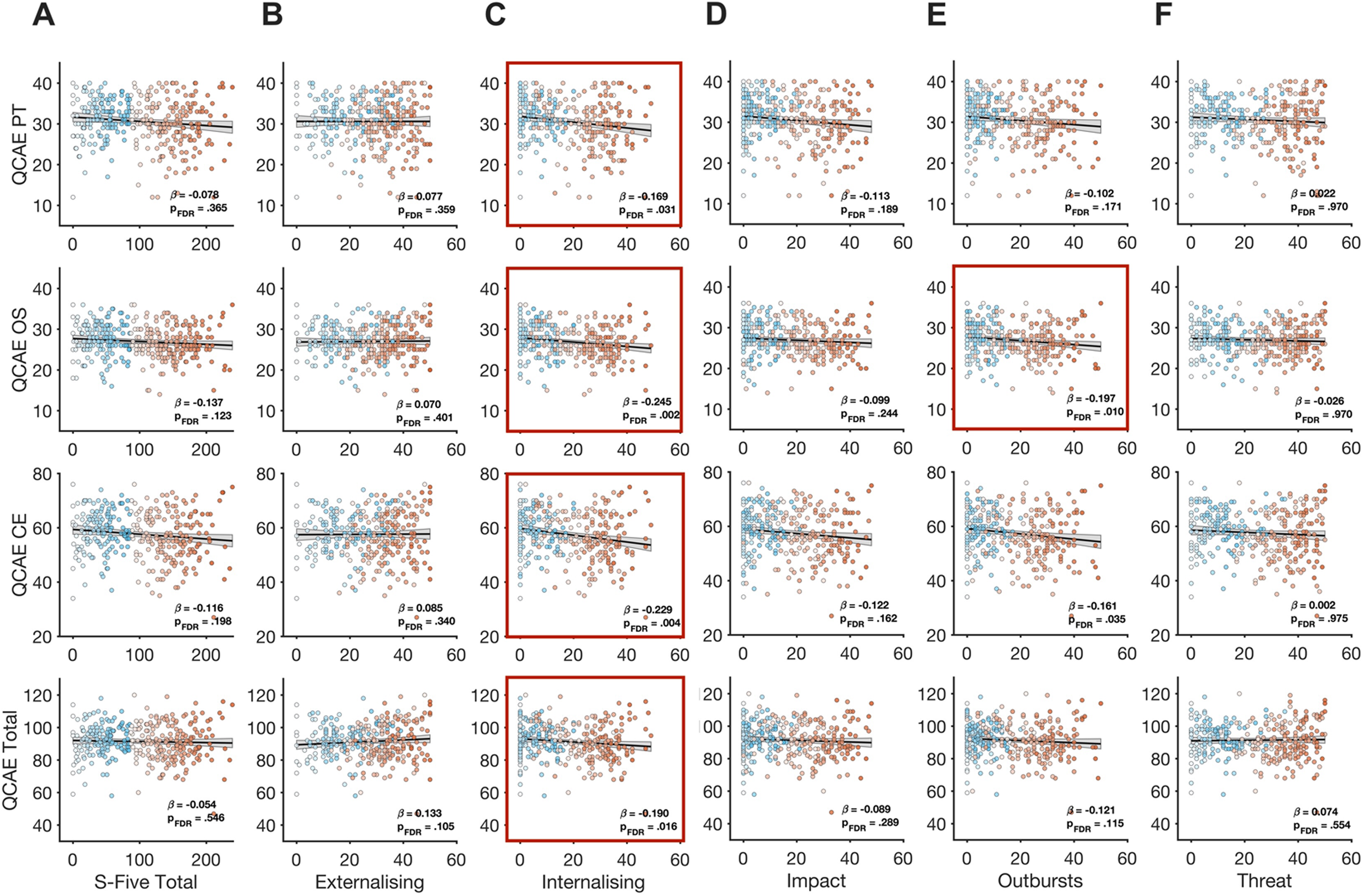

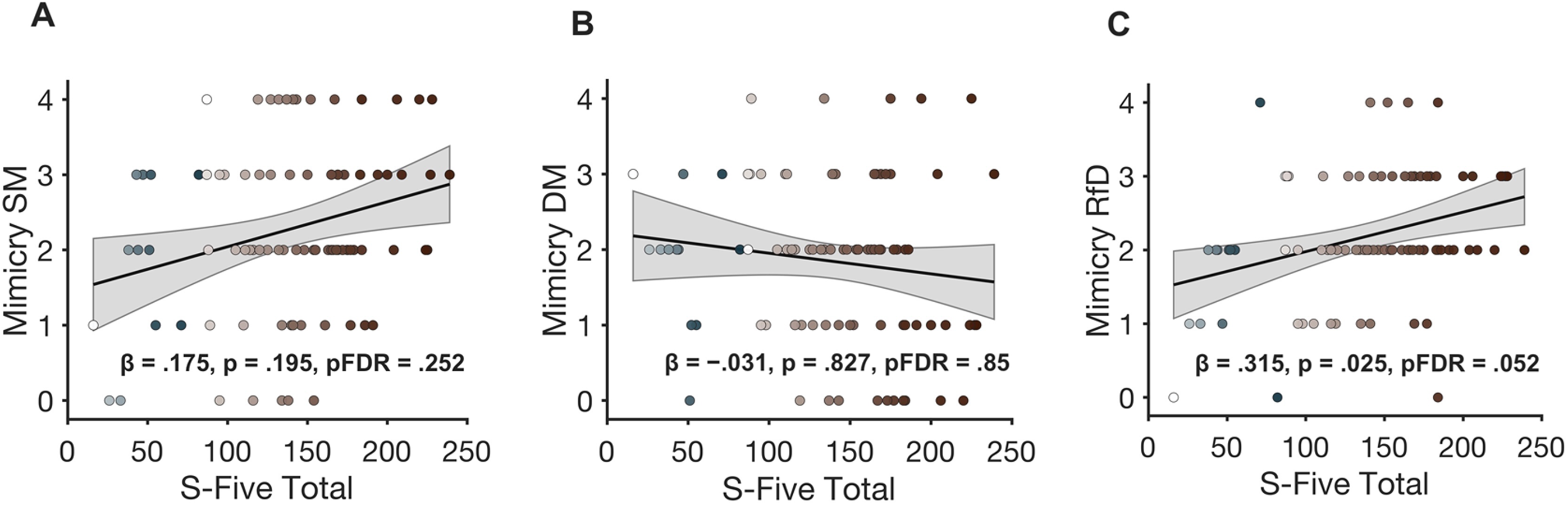

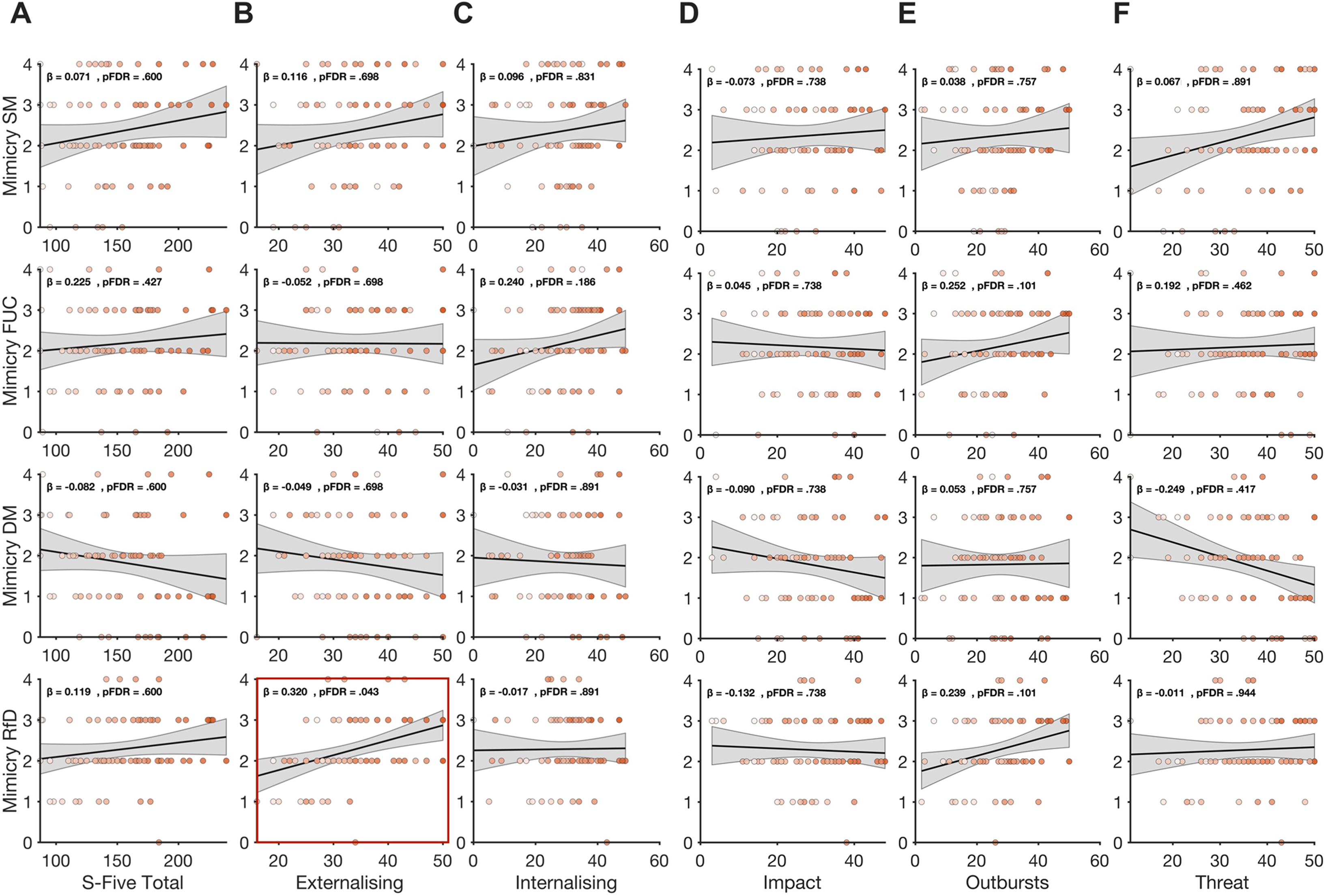

